# Long-term methylome changes after experimental seed demethylation and their interaction with recurrent water stress in *Erodium cicutarium* (Geraniaceae)

**DOI:** 10.1101/2023.01.19.524693

**Authors:** Francisco Balao, Mónica Medrano, Pilar Bazaga, Ovidiu Paun, Conchita Alonso

**Affiliations:** Departamento de Biología Vegetal y Ecología, Universidad de Sevilla, Apdo. 1095, 41080 Sevilla, Spain; Estación Biológica de Doñana, CSIC, Avenida Américo Vespucio 26, 41092 Sevilla, Spain; Department of Botany and Biodiversity Research, University of Vienna, 1030 Vienna, Austria

**Keywords:** abiotic stress, bisulfite sequencing, BsRADseq, differential methylation, DNA methylation, drought, epigenetics, 5-Azacytidine

## Abstract

- The frequency and length of drought periods are increasing in subtropical and temperate regions worldwide. Epigenetic responses to water stress could be key for plant resilience to this largely unpredictable challenge. Experimental DNA demethylation together with application of a stress factor stands as a suitable strategy to uncover the contribution of epigenetics to plant responses to stress.
- We analysed leaf cytosine methylation changes in adult plants of the Mediterranean weed, *Erodium cicutarium*, after seed demethylation with 5-Azacytidine and/or recurrent water stress in a greenhouse. We used bisulfite RADseq (BsRADseq) and a newly reported reference genome for *E. cicutarium* to characterize methylation changes in a 2×2 factorial design, controlling for plant relatedness.
- In the long-term, 5-Azacytidine treatment alone caused both hypo and hyper-methylation at individual cytosines, with substantial hypomethylation in CG contexts. In control conditions, drought resulted in a decrease in methylation level in all but CHH contexts. In contrast, the genome of plants that experienced recurrent water stress and had been treated with 5-Azacytidine increased DNA methylation level by ca. 5%.
- Seed demethylation and recurrent drought exhibited a highly significant interaction in terms of global and context-specific cytosine methylation supporting an epigenetic contribution in response to stress at molecular level.

## Introduction

Climate is changing. The period from 1983 to 2012 was likely the warmest 30-year period of the last 1400 years in the Northern Hemisphere (IPCC, 2014). Strong regional and temporal variability and an increased frequency of extreme events appear linked to global warming, together with a latitudinal redistribution of precipitation leading to increased flooding and drought events on different regions worldwide (Zhang *et al*., 2007). The Mediterranean Basin is an especially vulnerable region to predicted global changes encompassing increase of summer temperature, reduction of winter precipitation and more frequent heat waves and drought events (Giorgi & Lionello, 2008). Plants in this region are generally used to inconsistent precipitation and to drought (Cowling *et al*., 2005; Cook *et al*., 2016) and could, thus, exhibit increased resilience to such stress factor (Matesanz & Valladares, 2014; Balao *et al*., 2018; López□ Rubio *et al*., 2022). Analyzing the mechanisms behind adaptability to recurrent drought could thus be key to understand potential plant responses to the predicted effects of climate change.

Mounting experimental evidence substantiates that epigenetic mechanisms are involved in plant responses to stress (e.g., Gutzat & Mittelsten Scheid, 2012; Sun *et al*., 2020). A comprehensive understanding of the interrelationships between chromatin configuration, DNA methylation, histone tail modification and the activity of transposable elements in response to stress is being acquired for model and crop species with well-annotated reference genomes (Mirouze & Paszkowski, 2011; Gutzat & Mittelsten Scheid, 2012; Bäurle, 2018). On the other hand, research on non-model plants has largely focused on changes in cytosine methylation, providing evidence this is a key element of plant epigenomes (Niederhuth & Schmitz, 2017). In plants, methylation can be found at CG, CHG and CHH contexts (where H = A, C or T) because of the action of different families of methyltransferases which can be independently regulated (Lyko, 2018). Experimental studies provide support for a relevant role of DNA methylation in response to several abiotic stressors and particularly to water deficit and salt excess (Peng & Zhang, 2009; Alonso *et al*., 2016; Banerjee & Roychoudhury, 2017). In addition, experimental modification of DNA methylation profiles by applying inhibitors of DNA methyltransferases (DNMTs) such as Zebularine and 5-Azacytidine (Lopez *et al*., 2016) has further substantiated the potential epigenetic regulation behind phenotypic responses to stress in non-model plants (Verhoeven *et al*., 2010; Herman *et al*., 2016; Rendina González *et al*., 2016). Understanding the stability of epigenomic changes induced by artificial demethylation at seed stage could be particularly relevant to better evaluate the ability to use epigenetic information and seed priming for crop improvement (Gallusci *et al*., 2017; Springer & Schmitz, 2017).

Moving forward from anonymous markers and indirect evidence of epigenetic responses to environmental stress in non-model species is timely to better understand the ecological relevance of epigenetic variation in natural plant populations (Richards *et al*., 2017). Intrinsic plant (epi)genomic features (Springer *et al*., 2016), life-history traits (Verhoeven & Preite, 2014) and the extrinsic factors associated to the environment in which species have evolved (Balao *et al*., 2018; Herrera *et al*., 2019) might condition the magnitude of epigenetic changes in response to specific stress factors and, thus, generalizations from results obtained for model and crop species is not straightforward (Richards *et al*., 2017). Restriction site-associated DNA sequencing (RADseq hereafter) has become a popular reduced representation method for non-model organisms. RADseq uses restriction enzymes to guide complexity reduction to sequencing only a representative fraction of the genome, and can be implemented with or without prior genomic resources (Andrews *et al*., 2016). Similarly, reduced representation bisulfite sequencing (RRBS) methods provide a cost-effective alternative to Whole Genome Bisulfite Sequencing (WGBS) for the analysis of DNA methylation differences between groups of samples in species that lack a well-annotated genome (Paun *et al*., 2019). RRBS provide insights into the magnitude and the genomic location of substantial methylation changes, a relevant step forward because the effects of DNA methylation on gene expression, transposon activation and specific phenotypic traits depend on both the sequence context and genomic location (Hirsch *et al*., 2012; Niederhuth & Schmitz, 2017).

In this paper, we use BsRADseq (Trucchi *et al*., 2016) to investigate changes in DNA cytosine methylation in leaves of adult individuals of the annual herb *Erodium cicutarium*, after experimental seed demethylation and recurrent drought along their life-cycle. Addition of inhibitors of DNA methyltransferases (DNMTs) at seed germination has been proofed to be effective in reducing global cytosine methylation in seedlings of several species including this one (Alonso *et al*., 2017). However, a detailed analysis of the genome-wide effects of such demethylating agents is only available for the model species *Arabidopsis thaliana*, which suggests a significant dose dependent reduction in all cytosine contexts in 10 d seedlings (Griffin *et al*., 2016). To the best of our knowledge no study has applied genomic tools to assess if these early effects are maintained later in the life cycle, although repeated foliar spraying and foliar injection of DNMT inhibitors have been proposed as alternative methods to continuously alter DNA methylation profiles along plant life-cycle (see Puy *et al*., 2018; Herrera *et al*., 2019, respectively). Thus, evaluating the magnitude and specific patterns of demethylation induced after seed treatment and their stability up to the reproductive stage, is a critical point for assessing the utility of this method to reveal the associations between DNA methylation and plant phenotypic variance, and its contribution to stress response in a broad range of plant species (Richards *et al*., 2017; Alonso *et al*., 2019b) as well as its potential relevance for epigenetic priming of seed crops (Gallusci *et al*., 2017). In particular, only long-lasting demethylation effects that persist until the adult reproductive stage, enduring pollen maturation and seed formation, might be relevant for transgenerational transmission of induced epigenetic variants.

Stress exposure may prime plant individuals and/or their offspring for improved response after similar or other stress events (Walter *et al*., 2013; Pandey *et al*., 2015; Dangi *et al*., 2018). However, enduring responses could also compromise plant fitness under different conditions (Tricker, 2015; Crisp *et al*., 2016; Latzel *et al*., 2016; Douma *et al*., 2017). In this study we focused on recurrent drought because it is characteristic of the Mediterranean ecosystems in which *E. cicutarium* lives and also because we were uncertain of the time lapse for eliciting strain and subsequent recovery after a single extreme drought event (see e.g., Walter *et al*., 2011; López-Jurado *et al*., 2016). Our specific questions were: do genomes of adult plant leaves exhibit any signature of DNMT inhibition by 5-Azacytidine at seed stage? which are the genomic methylation changes associated to recurrent water stress along individual *E. cicutarium* life cycle? is there any interaction between these two factors?

## Materials and Methods

### Study species

*Erodium cicutarium* (L.) L’Hér. (Geraniaceae), is a diploid annual herb native to Mediterranean Europe, North Africa and western Asia, that is currently distributed globally in temperate areas with hot summers (Fiz-Palacios *et al*., 2010; Francis *et al*., 2012). It has a fast-growing cycle and autonomous self-pollination that make it a good candidate for experimental studies in greenhouse.

### Experimental design

Demethylation and recurrent water stress were applied to the offspring of plants cultivated in the greenhouse. A first generation was grown in order to minimize the effects of heterogeneous growing conditions of maternal wild plants, which could affect their offspring phenotype and DNA methylation profiles (Latzel, 2015). The parental generation (F0) were adult plants at two *E. cicutarium* natural populations located in Cazorla mountains (Jaén province, SE Spain). Seeds were removed from fruits collected in the field, scarified, and germinated in universal substrate (COMPO SANA ^®^) mixed in 3:1 with perlite (substrate hereafter). The first generation (F1) seedlings were grown in 1L pots with the same substrate; pots were grouped in trays, watered twice per week, and trays were periodically rotated within the greenhouse (16h light; 25-20 °C) until the end of reproduction (ca. 6 months). Autonomously-pollinated fruits were collected in paper bags, and stored at room temperature (see Alonso *et al*., 2017 for further details.

Four months later, seeds were removed from fruits, weighed and slightly scarified with sandpaper. A demethylation treatment was applied to half of the seeds as described in Alonso *et al*. (2017). In brief, scarified seeds were submerged in 150 μl of either Control (water with DMSO 97:3, v:v) or a 0.5 mM solution of 5-Azacytidine (Sigma A2385-100mg) for 48 h at 4 °C. Immediately after, all seeds were individually transferred to seedling plug trays filled up with substrate. Trays were saturated in water and placed under greenhouse conditions (16h light; 25-20 °C). After 39 d of sowing, seedlings were transplanted to 1L pots. Extra seedlings were collected to confirm the effects of the demethylation treatment, which were detected as a lower global cytosine methylation percentage in leaf DNA and production of shorter and fewer leaves compared to controls at this developmental stage (Alonso *et al*., 2017).

Three days later, when seedlings were 6 weeks old, the recurrent water-stress (WS hereafter) treatment started. For each maternal line, half the offspring was watered at field capacity twice per week (i.e., receiving optimal watering; WW hereafter) and half the offspring was watered at field capacity once every 10-11 d until the end of the experiment (WS). Flowering started soon after transplantation and reached the peak (75 % individuals with flowers) at week 10. Demethylated plants tended to flower ca. five days later than their untreated relatives (C. Alonso and M. Medrano unpub. data). The experiment finished when plants were 16 weeks old, and some plants started showing signs of senescence.

### Sample processing

Methylation analyses were conducted on DNA extracted from leaves of a subset of reproductive F_2_ individuals according to the design: 2 provenances * 4 F_1_-mothers * 2 demethylation levels * 2 watering levels = 32 F2-individuals). A sample of 2-3 full grown leaves without signs of damage or senescence of each F2 individual was collected, placed in labeled paper bags and dried at ambient temperature in sealed containers with abundant silica gel.

Dried samples were homogenized to a fine powder using a Retsch MM 200 mill. Total genomic DNA was extracted using Bioline ISOLATE II Plant DNA Kit and quantified using a Qubit fluorometer 2.0 (Thermo Fisher Scientific, Waltham, MA, USA).

### RADseq and BsRADseq library preparation

The RADseq and BsRADseq libraries were prepared following the laboratory protocol detailed in Trucchi *et al*. (2016), with minor changes. In particular, we modified the restriction enzyme used and employed the more frequent cutter *PstI* (6 bp restriction site), in order to maximize the density of markers for differential methylation analysis.

We used 20 units of *Pst*I HF restriction enzyme (New England Biolabs) to digest 800 ng DNA per individual and ligated 100 nM P1 barcoded methylated adapters over night at 16°C. Groups of samples barcoded with different P1 barcodes were pooled, sheared to a target peak of 400 bp (with a Covaris E220 focused-ultrasonicator) and ligated to methylated P2 adapters. We employed a set of methylated P1 and P2 barcoded adapters to protect their sequence from modification during bisulfite treatment. The fragments were constructed with double barcoding system: eight different 5 bp barcodes were inserted with the P1 adaptor in combination with four different P2 adaptor with 6 bp long barcodes. Our barcodes differed in at least four bases from each other. At this step, one aliquot was separated and directly amplified by PCR as a standard RADseq library. For the other aliquot, bisulfite conversion of non-methylated Cs into Us was performed after P2 adapter ligation by treating the library with MethylEdge Bisulfite Conversion System kit (Promega). PCR amplification was executed using KAPA HiFi HotStart Uracil+ MasterMix (Kapa) for 23 cycles to convert all Us into Ts. The final two libraries were sequenced at the VBCF Vienna (https://www.viennabiocenter.org/vbcf/next-generation-sequencing/) as paired-end 125 bp reads in two lanes of an Illumina HiSeq 2500 machine.

### Assembly of Erodium cicutarium draft genome

*Erodium cicutarium* has a genome size of 1C = 1.20 pg and CG content = 0.4 (Pustahija *et al*., 2013). A draft genome for *E. cicutarium* was assembled from DNA extracted from a single individual reared after two events of autonomous selfing under greenhouse conditions with a paired-end strategy in Illumina HiSeq X (PE150) at AllGenetics (www.allgenetics.eu). KmerGenie 1.7038 (Chikhi & Medvedev, 2014) was used to estimate the best kmer in the pair reads and k = 71 was selected as the start point for de novo assemble with ABySS 2.02 (Simpson *et al*., 2009). Plastid and mitochondrion genomes were handled separately. Reads were mapped back to the assembled output to determine coverage depth using BWA 0.7.12 (Li & Durbin, 2010). SAMtools 0.1.19 (Li *et al*., 2009) was used to remove bad quality and secondary alignments. To avoid possible inconsistencies or the presence of contaminant sequences, contigs/scaffolds with a mean coverage < 10x were discarded, as well as those shorter than 1,000 bp.

RepeatModeler 1.0.11 and RepeatMasker 4.0.7 were used to identify and mask the predicted repeats in the filtered assembly. Geneid 1.4 (Alioto *et al*., 2018) and SMA3s (Muñoz-Mérida *et al*., 2014) were used to perform the prediction of genes and their subsequent functional annotation. The manually annotated protein database Swiss-Prot (taxonomic division: plants) available in the UniProt database (The Uniprot Consortium, 2007) was used as reference. In addition, the predicted genes were also compared against GenBank’s nr protein database including only the Embryophyta taxa entries by using a BLAST+ 2.6.0 (Camacho *et al*., 2009).

### RADseq and BsRADseq data processing and alignment

Barcoded raw Illumina reads were processed with deML (Renaud *et al*., 2015) and STACKS v.2.0Beta8 (Rochette *et al*., 2019). To demultiplex the individuals and remove low quality data, the program *process radtags* was run with the following settings: PstI as restriction enzyme, restriction site check at the beginning of the reads disabled (due to BS conversion affecting the sequence of the restriction site), discarding reads with low quality scores according to default parameters, and rescuing barcodes and RADtags with two sequencing error (following Trucchi *et al*., 2016). The paired-end fastq files obtained were filtered to retain only full-length reads (i.e., P1 120 bp and P2 125 bp after barcode trimming), with no adaptor contamination, and unambiguous barcode sites.

The standard RADseq sequences of each of the 32 samples were mapped to the reference genome (see below) to identify variable sites (i.e., SNPs) involving a cytosine in the reference sequence. C/T polymorphisms were subsequently filtered and masked with vcftools for subsequent methylation analyses (see Trucchi *et al*., 2016 for details). BsRADseq reads of each of the 32 samples were mapped to the draft reference genome using the mapping routine available in *Bismark* (Krueger & Andrews, 2011) specifically designed for dealing with bisulfite converted reads with the core aligner Bowtie2 (Langmead & Salzberg, 2012) in a non-directional modus and allowing up to four mismatches for a 120 bp read (options: --non_directional -L 32 -D 10 -R 1 --score_min L,0,-0.2). The sodium bisulfite conversion efficiency rates were assessed by calculating cytosine methylation levels in the chloroplast genome (Schmidt *et al*., 2017) and were found globally satisfactory with conversion ≥ 99.2% in all cases (Table S1). After mapping, we checked the summary report for each individual, recording the mapping efficiency, the number of cytosines screened, the distribution among the different contexts (CG, CHG, and CHH) and the differential representation of original strands versus complementary to original strands. As expected, the reads mainly mapped (> 99%) complementary to either the top or the bottom strand of the reference genome due to peculiarities of the bsRADseq protocol (Trucchi *et al*., 2016) and the stochastic orientation of the contigs in the reference genome. At supplementary materials, Table S1 presents a summary of sequencing and mapping results for each individual sample. The next step was to extract the methylation information of each cytosine position using the *Bismark_methylation_extractor* routine ignoring the first 4 bp in the reads, including the “–no_overlap” flag to prevent counting the same cytosine if covered by both the forward and reverse read and producing cytosine reports (CX_report files in the Bismark output) for each individual sample with information on all cytosine contexts and all strands. Output was then merged by sequence context (CG, CHG, CHH) for downstream analysis.

### Global methylation analyses

Differential methylation analyses were conducted in R software 4.2.1 (R Core Team, 2022). Firstly, the global effects of WS and 5-Azacytidine agent (5-Aza hereafter) on the genome wide cytosine methylation level were evaluated using a generalized linear model (GLM) with binomial error and logit link. The dependent variable was the proportion of methylated Cs to unmethylated Cs in each sample (i.e., independent of their genomic position) and the independent ones were WS and 5-Aza and their interaction effect (5-Aza + WS), as well as the F_1_-mother identity (as proxy of genotype), all considered as fixed factors. The 3-way interaction was not assessed due to absence of maternal replicates within each factor combination level, still the experimental design controlled for any potential bias that maternal identity could introduce by always including one plant from each maternal family in the four factorial combinations. The analysis was run for the whole dataset and separately for each sequence context (CG, CHG, CHH).

### Differential methylation analyses

Furthermore, to gain insight into the specific effects of 5-Aza and WS on the methylation patterns with an explicit genomic context, differentially methylated cytosines (DMCs) at CG, CHG and CHH sites were estimated using the callDML command in DSS (dispersion shrinkage for sequencing data) package v.2.34.0 (Feng *et al*., 2014) with a 2 x 2 ANOVA design including the effects of WS, 5-Aza and their interaction (DMLfit. multiFactor = WS + 5-Aza + WS:5-Aza). In brief, DSS allows complex experimental designs (including interaction effects), based on a beta-binomial regression model with arcsine link function, and it takes advantage of using a shrinkage estimator of the dispersion parameter based on a Bayesian hierarchical model to reduce the dependence of variance on mean. P-values were adjusted for multiple testing using Benjamini and Hochberg false discovery rate (FDR) procedure. For each effect (WS, 5-Aza and their interaction), DMCs with a corrected q-value < 0.05 and > 10% difference in the methylation level were considered candidate DMCs.

To investigate the methylation changes of the DMCs with significant interaction effects, we used the optimized K-mean clustering minimizing the sum-of-squares within cluster (WCSS; Witten & Tibshirani, 2010) on the average methylation of each of the four levels of treatment combination (Control, 5-Aza, WS, 5-Aza + WS). Methylation of DMCs and clustered DMCs were visualized using a heatmap created with the R package *pheatmap* v. 1.0.12 (Kolde, 2019).

### Annotation of DMC-associated genes

DMC-associated genes were defined as genes with DMCs within the gene body region (coding sequence region, CDS, and introns) and/or promoter regions (1 kb upstream the CDS). In addition, DMCs located within regions annotated as transposable elements were separately analyzed.

## Results

### Draft genome assembly and annotation

A total of 225 million pairs of reads have been used in the assembly giving a mean coverage of 33.8x (SD = 33.5). The assembly of k = 97 was chosen as the best reconstruction. A total of 65,050 contigs/scaffolds were kept in the final assembly (total length 628,531,192 bp; N50 = 12,445; L50 = 15,770; GC% = 41.0). The BUSCO 3.0 (Waterhouse *et al*., 2018) comparison to Embryophyta gene set yielded 75.5 % complete, 8.5 % fragmented and 15.9 % missing BUSCO orthologs. SMA3s was able to annotate a total of 43,537 genes with 99.9 % associated to a specific GO term. Furthermore, a total of 2,135 repeat families were predicted by RepeatModeler and 38.5 % of the genome was masked as repetitive.

### BsRADseq library output

We obtained between 2,654,652 and 31,840,142 pairs of reads for the individual samples. After quality filtering, between 1,133,391 and 11,417,213 paired sequences per individual were retained (Table S1), corresponding to 35.8% - 42.7% of the raw reads. On average, the number of pair reads per sample was 4,486,477, the GC content was 28.9 % (± 1.3), a low figure, as expected for bisulfite treated samples. The mapping efficiency to our draft reference genome averaged 38.4 %, and ranged from 35 % to 42.5 % across study individuals, resulting in an average coverage of 98.6x +/- 2.1x (Table S1).

### Global DNA methylation changes

The global cytosine methylation level in the genomes of adult *E. cicutarium* leaves, as estimated by *Bismark*, varied widely across study plants and conditions, ranging from 16.7% to 30.6% (Table 1), and averaged 22.8 % in the genomes of untreated plants (i.e., controls). Furthermore, the average cytosine methylation level was different at the three sequence contexts analyzed, in CG context methylation was estimated as 66.0%, whereas 30.8% of cytosines were methylated in CHG and only 9.3% in CHH contexts (Table 1). CG sites were the rarest in the genome, but because of their high mean methylation level, they represented a major proportion of methylated cytosines, contributing the most to the overall cytosine methylation that globally averaged 23.2 % (SD 3.7) in our full set of samples.

**Table 1.** Genome-wide methylation levels as total and for each sequence context for the 32 samples. C, control; A, adult plants obtained after 5-Aza treatment of seeds; WW, optimal watering; WS, water stress.

| Treatments |  | Family | Sample | No. | mC | mCG | mCHG | mCHH |
| --- | --- | --- | --- | --- | --- | --- | --- | --- |
| 5-Aza | Watering |  |  |  |  |  |  |  |
|  |  |  |  | cytosines | (%) | (%) | (%) | (%) |
| C | WW | PT_1031 | CC_253 | 92353730 | 18.4 | 58.6 | 22.3 | 6.3 |
| C | WW | PT_1001 | CC_225 | 108547644 | 23.3 | 67.7 | 32.8 | 8.4 |
| C | WW | PT_0969 | CC_147 | 47011293 | 18.7 | 60.2 | 21.9 | 7.4 |
| C | WW | PT_0925 | CC_001 | 124533805 | 24.5 | 67.5 | 32.9 | 10.5 |
| C | WW | CH_1307 | CC_491 | 39099076 | 25.5 | 68.8 | 34.1 | 11.4 |
| C | WW | CH_1245 | CC_433 | 22052707 | 23.2 | 66.2 | 31.9 | 9.8 |
| C | WW | CH_1233 | CC_409 | 94508383 | 17.1 | 58.5 | 18.4 | 6.6 |
| C | WW | CH_1187 | CC_291 | 85841474 | 25.9 | 69.5 | 35.8 | 10.6 |
| C | WS | PT_1031 | CD_259 | 128071349 | 17.8 | 58.6 | 20.4 | 6.4 |
| C | WS | PT_1001 | CD_219 | 116121937 | 27.8 | 69.2 | 43.5 | 9.8 |
| C | WS | PT_0969 | CD_163 | 128356306 | 23.0 | 65.4 | 30.3 | 9.5 |
| C | WS | PT_0925 | CD_005 | 213718116 | 21.8 | 63.1 | 25.8 | 9.7 |
| C | WS | CH_1307 | CD_485 | 22299814 | 23.7 | 69.4 | 29.0 | 10.2 |
| C | WS | CH_1245 | CD_445 | 29859487 | 21.4 | 63.3 | 27.3 | 8.7 |
| C | WS | CH_1233 | CD_431 | 87893575 | 16.7 | 58.3 | 17.5 | 5.9 |
| C | WS | CH_1187 | CD_297 | 50225708 | 25.8 | 71.8 | 35.8 | 10.0 |
| A | WW | PT_1031 | AC_252 | 164753131 | 24.9 | 68.8 | 34.5 | 9.3 |
| A | WW | PT_1001 | AC_220 | 97316260 | 18.9 | 57.8 | 26 | 6.4 |
| A | WW | PT_0969 | AC_146 | 46070213 | 19.0 | 61.2 | 22.9 | 6.6 |
| A | WW | PT_0925 | AC_016 | 75973105 | 27.6 | 70.4 | 37.6 | 11.9 |
| A | WW | CH_1307 | AC_494 | 44354076 | 24.4 | 69.7 | 31.6 | 10.3 |
| A | WW | CH_1245 | AC_438 | 21496364 | 25.7 | 69.6 | 35.6 | 10.7 |
| A | WW | CH_1233 | AC_422 | 99564473 | 21.2 | 63.0 | 26.3 | 8.9 |
| A | WW | CH_1187 | AC_300 | 54846691 | 18.1 | 59.4 | 21.2 | 6.8 |
| A | WS | PT_1031 | AD_254 | 137114375 | 25.5 | 67.2 | 37.4 | 9.4 |
| A | WS | PT_1001 | AD_222 | 38996631 | 25.7 | 67.9 | 37.9 | 8.8 |
| A | WS | PT_0969 | AD_164 | 85184218 | 24.5 | 68.9 | 33.3 | 9.3 |
| A | WS | PT_0925 | AD_020 | 105676423 | 28.8 | 72.6 | 40 | 13.2 |
| A | WS | CH_1307 | AD_482 | 38691859 | 22.1 | 65.7 | 28.0 | 9.1 |
| A | WS | CH_1245 | AD_436 | 34188955 | 28.1 | 73.4 | 39.1 | 12.3 |
| A | WS | CH_1233 | AD_416 | 82021987 | 23.6 | 64.9 | 30.1 | 10.2 |
| A | WS | CH_1187 | AD_292 | 74545405 | 30.6 | 76.0 | 44.0 | 13.0 |

The wide range of variance in global methylation level across individuals was congruent with the significant effect of the F1-mother factor (as proxy of genotype) on the proportion of methylated Cs (LR-χ^2^= 5421658, P < 0.0001, Fig. S1) and the impact of the two treatments, whose effects exhibited a significant interaction on this variable (LR-χ2= 1036036, P < 0.0001. Whereas the 5-Aza seed exposure weakly affected the Cs methylation in leaves of adult plants grown under optimal watering conditions, a global and significant increment in methylation level (~5%) was observed in the genome of leaves of plants that experienced recurrent WS and had been treated with 5-Aza at seed stage (Fig. 1). Changes in methylation level of Cs at the three sequence contexts followed similar patterns (Fig. 1). Interestingly, whereas drought stress appeared to decrease the global methylation level and that at CG and CHG contexts, it increased DNA methylation at CHH sites that tend to be associated with transposable elements (e.g., Martin *et al*., 2021).

**Fig. 1.**
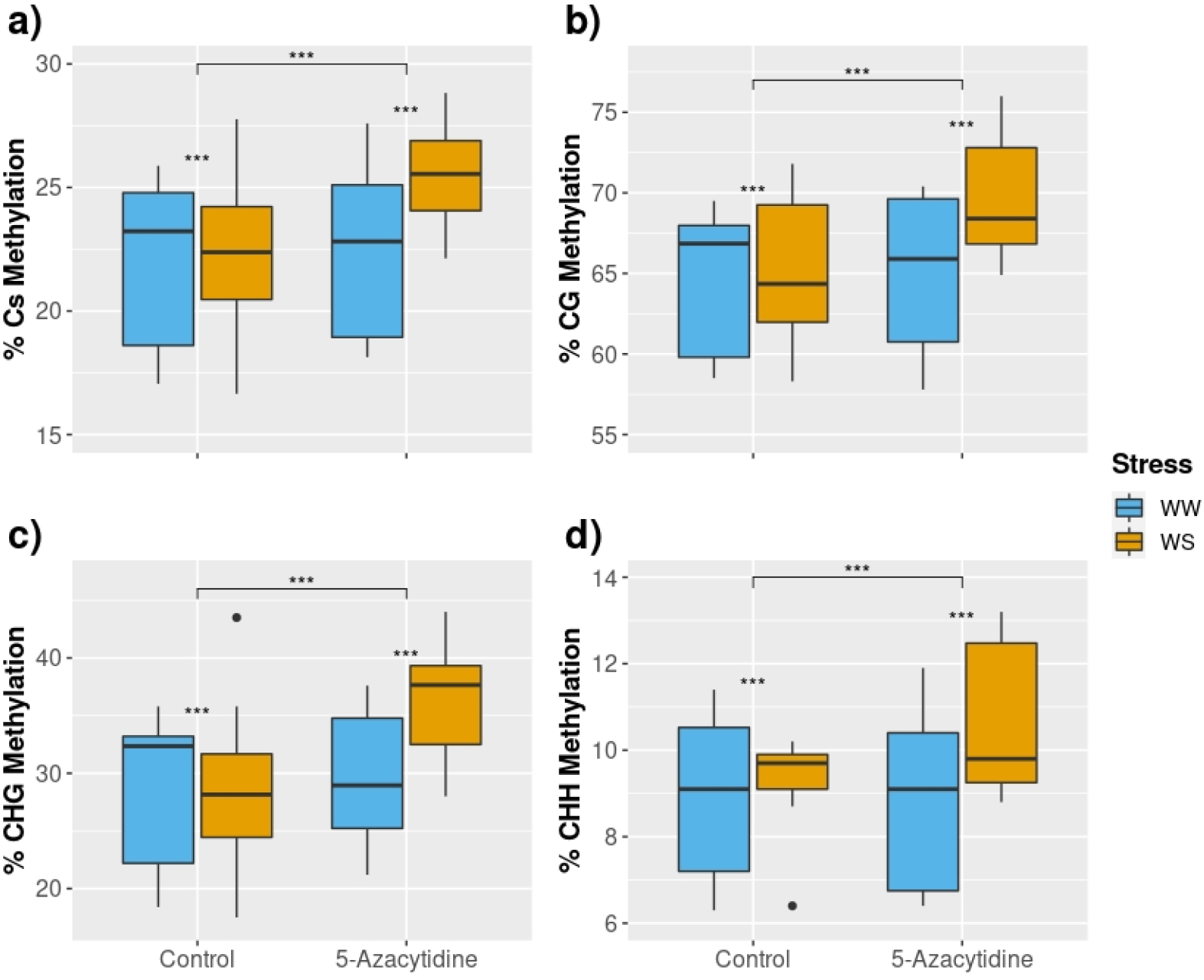
Boxplots of the percentage of methylated cytosines found on leaf genomes of adult plants obtained from seeds with no treatment (Control) and seeds treated with 5-Azacytidine (5-Azacytidine), and grown with optimal watering (blue, WW) or recurrent water stress (yellow, WS). (a) Global methylation, (b) methylation percentage at CG, (c) methylation percentage at CHG, (d) methylation percentage at CHH. *** P<0.001. N=8 for each factor combination level. Note that in all cases the interaction between the two study factors was statistically significant (see Results for further details).

### Genomic methylation changes induced by 5-Aza and recurrent water stress in leaves of E. cicutarium

A total of 3.1% of the 222,835 analyzed cytosine positions were identified as differentially methylated cytosines (DMCs) between control plants and those experiencing just 5-Aza, just WS or both treatments. An uneven distribution of DMCs was observed along the genome, the number of DMCs per scaffold differed from a null distribution (Wilcoxon’ test P < 0.001, Fig. S2), mainly because the largest fraction of scaffolds only got one or two DMCs regardless of their size and Cs abundance. Further, the frequency of DMCs in the three sequence contexts did not resemble their relative abundance, with the highest proportion being located in CHH (48.67%), followed by CG (33.31%), and the lowest being in CHG (18.02%).

If we look at the changes induced by the two treatments separately, seed exposure to 5-Aza had broader impact on the adult leaf methylome in terms of positions modified (3,897 DMCs) than the WS treatment (1,417 DMCs). This trend was maintained across the three sequence contexts (Fig. 2a) with 1,323 vs. 501 DMCs in CG context, 710 vs 235 in CHG and 1,864 vs 681 in CHH for 5-Aza and WS treatments, respectively.

**Fig. 2.**
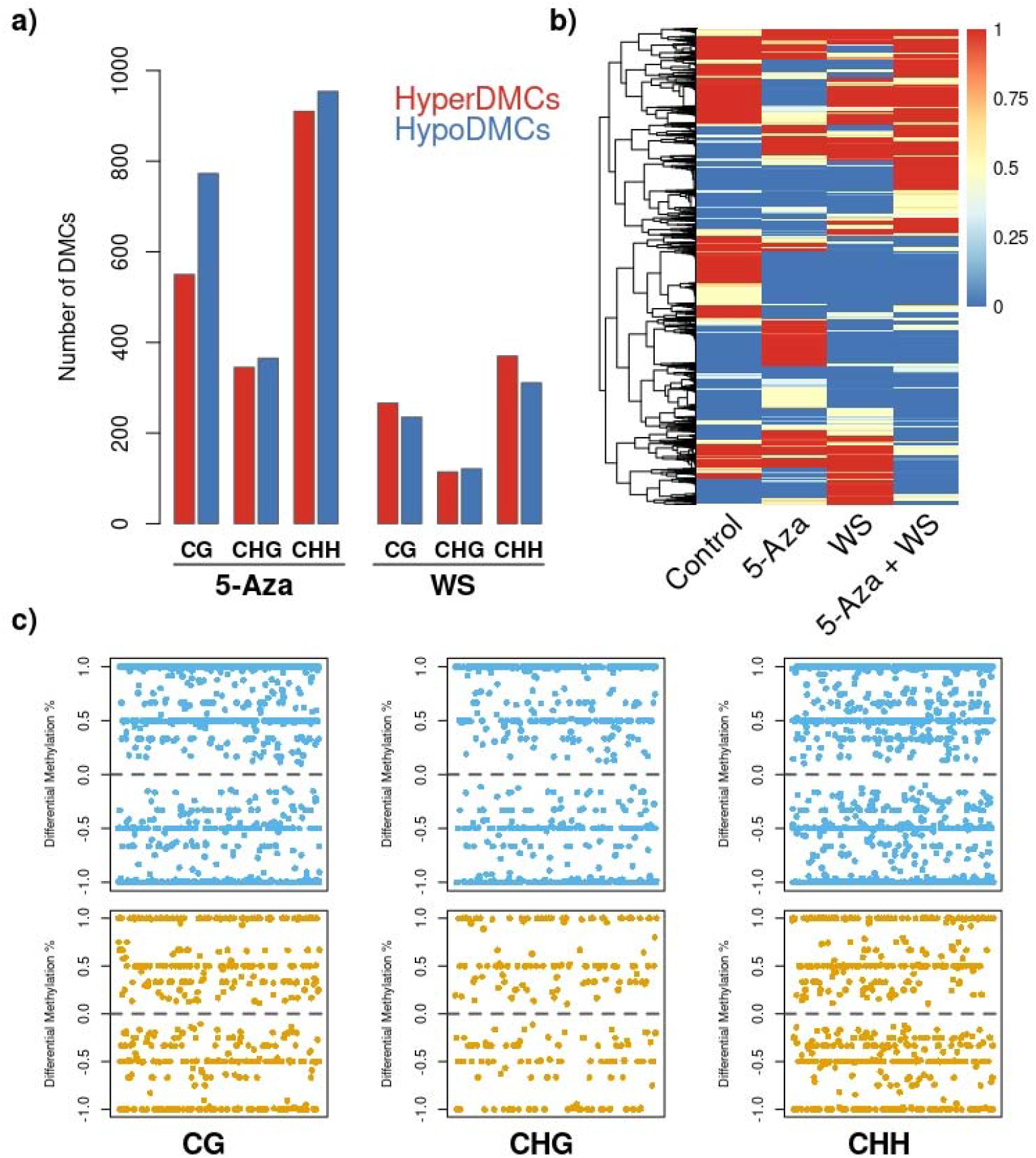
a) Number of Differentially Methylated Cystosines (DMCs) by context induced by application of 5-Azacytidine at seed stage (5-Aza) and/or recurrent water stress (WS) that were hyper- or hypo-methylated compared to control plants. b) Heat map visualization of the 6848 DMCs obtained. Average methylation level for each experimental group (8 samples each) is shown. DMCs are clustered using the complete linkage clustering, and the scale is shown on the right, in which red and blue correspond to a higher and a lower averaged methylation status, respectively. c) The lower panels show the methylation difference between the two conditions of each study factor, blue for 5-Aza and orange for WS, at every DMC evaluated on each of the three sequence contexts analyzed. Each dot represents a certain DMC position for which a Y-axis value of 1 indicates complete lack of methylation in treated plants and full methylation in control condition.

As regards the direction of the observed changes, rather unexpectedly, seed treatment with 5-Aza caused both hypo and hyperDMCs in adult leaves (Fig. 2). The number of hypoDMCs (in comparison to control) was higher (773) to the number of hyperDMCs (550) in the CG context, as expected by its predicted negative impact on the activity of the methylase enzyme MET1, whereas the number of positions altered in the two directions was similar at both CHG and CHH contexts (Fig. 2a). Furthermore, WS caused similar number of hypo and hyperDMCs at CG, CHG and CHH contexts (Fig. 2b-c).

### Synergistic, antagonistic and transgressive effects of combined 5-Azacytidine and water stress on leaf methylome

A total of 4,073 DMCs showed significant interactive effects between 5-Aza and WS treatments. Most DMCs were located in CHH (48.3%), followed by CG (33.6%), whereas the lowest proportion was identified in CHG context (18.1%). We clustered these DMCs by the average methylation level across treatments highlighting groups with distinct interaction patterns (Fig. 3a). The number of different patterns (i.e. k-means groups) was eight (which minimized the within group sum of squares, Fig. S3). The eight patterns could be resumed into two general trends (see dendrogram in Fig. 3 a). The first one included DMCs with a very low methylation level in most of the treatments (Clusters 1, 2, 3 and 4), usually including the control treatment. This “low-methylation group” represented most of the DMCs analyzed with more than 600 DMC per cluster. The second group (Clusters 5, 6, 7 and 8) showed the opposite trend including DMCs with a high methylation level (~100%) in most of the treatments.

**Fig. 3.**
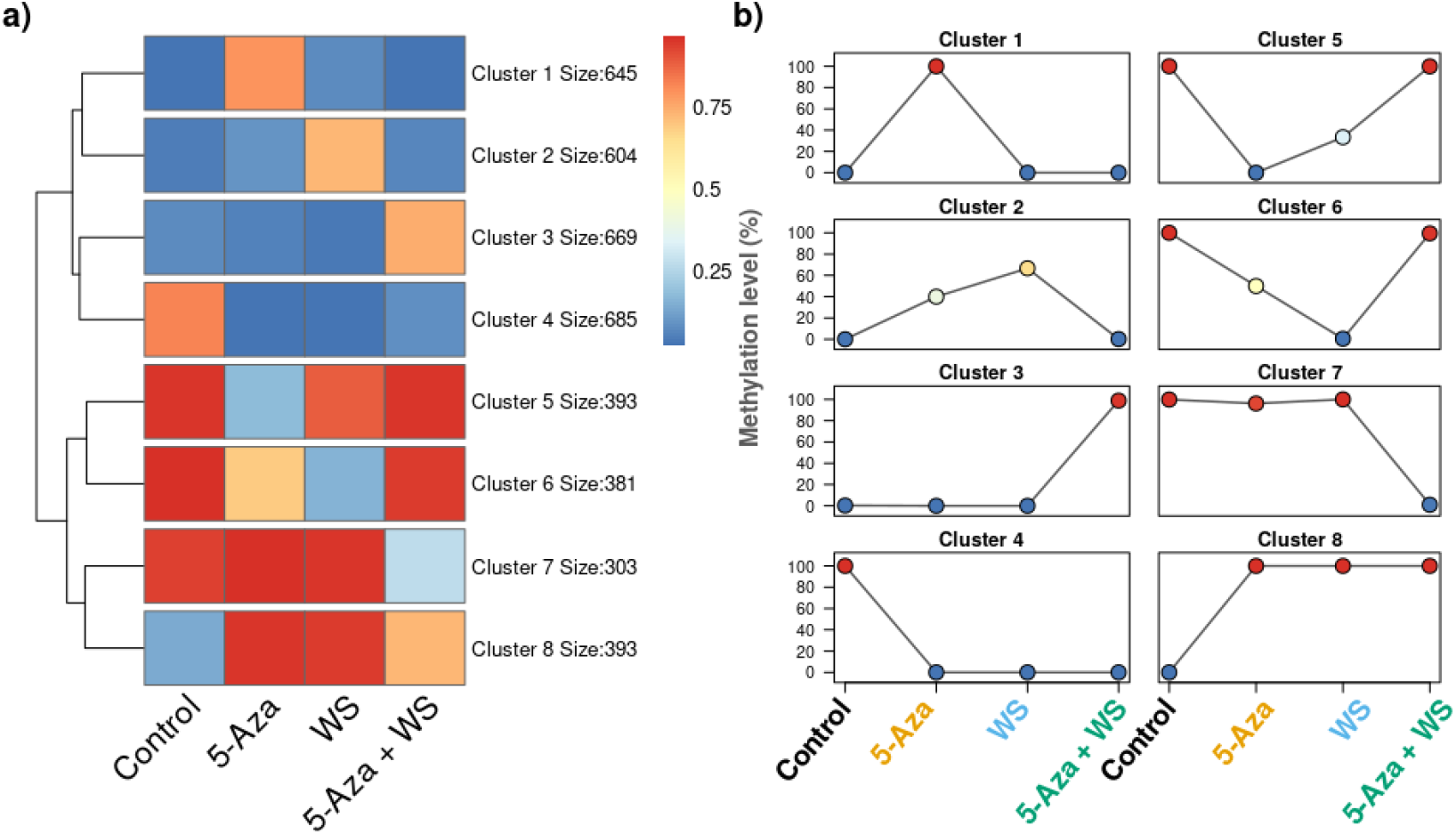
Visualization of 5-Azacytidine and water stress treatment interaction effects on the methylome. a) Hierarchical clustering of the 4073 DMCs whose methylation changes exhibited a significant interaction between the two study factors and heatmap of K-means clusters (K = 8) showing common DMC patterns. The number of cluster minimized the within cluster sum of squares. Size depicts the number of DMC within each cluster. b) Average methylation levels for each experimental group on each K-means cluster.

Furthermore, the k-mean clustering allowed us to unveil that many of the DMC were fully methylated or non-methylated in the genomes of control plants identifying hypo- and hypermethylation for the 5-Aza and WS treatments in comparison to the control (Fig. 3b, clusters 4 and 8 respectively). In addition, distinctive hyper- and hypomethylation was detected for 5-Aza + WS treatment (Fig. 3b, clusters 3 and 7, respectively), singular hypermethylation and hypomethylation for the 5-Aza treatment (clusters 5 and 1, respectively), increased hypermethylation (following methylation increment with 5-Azacitidine) for the WS treatment (cluster 2), and the opposite trend (i.e. increased demethylation) for the WS treatment (cluster 6). From these patterns, we could discern the synergistic effects of 5-Aza and WS on the methylation (clusters 2, 4, 6 and 8). The antagonist effects of both treatments were also shown in the clusters 1 and 5. Furthermore, transgressive effects for treatments combination were detected (clusters 2, 3, 5, 6 and 7).

### Annotation of genomic regions overlapping DMC

We further annotated the genomic regions that were found to be significantly differentially methylated (Fig. 4). For the 5-Aza treatment, 43% of the CG DMCs, 36% of the CHG DMCs and 38% of the CHH DMCs overlapped with gene regions (exons, introns and promoters). These proportions were slightly higher for the WS treatment, 46%, 41% and 41% for CG, CHG and CHH DMCs, respectively. Remarkably, the highest proportions of DMCs overlapping promoters and introns were found in the CHH context for both treatments. In addition, on average, 21.3% of DMCs overlapped transposable elements. Additionally, 36% of the DMCs overlapping TEs appeared inserted in genic regions (introns and promoters, Fig. S4). The percentage of TEs with DMCs (from total DMCs) inserted in genic regions increased from CG (4%) and CHG (6.5%) to CHH context (11.5%). Furthermore, both treatments, 5-Aza and WS, showed similar proportions of DMCs overlapping TE (without accounting those inserted in genic regions) for each different context (Fig. 4). The proportions TE-overlapped DMCs increased from GC context (8% in 5-Aza) to CHH context (19% in 5-Aza). In addition, although a large amount of TE-overlapped DMC was uncategorized, the proportion of TE classes significantly differed from the genome distribution (χ2 = 149.43; df = 6; p < 0.001). *Copia* elements containing DMCs (234) showed a higher proportion than expected (47.5% vs. 25.4%) whereas the LINE elements (61) showed a lower proportion (12.3% vs. 27.9%; Fig. 5).

**Fig. 4.**
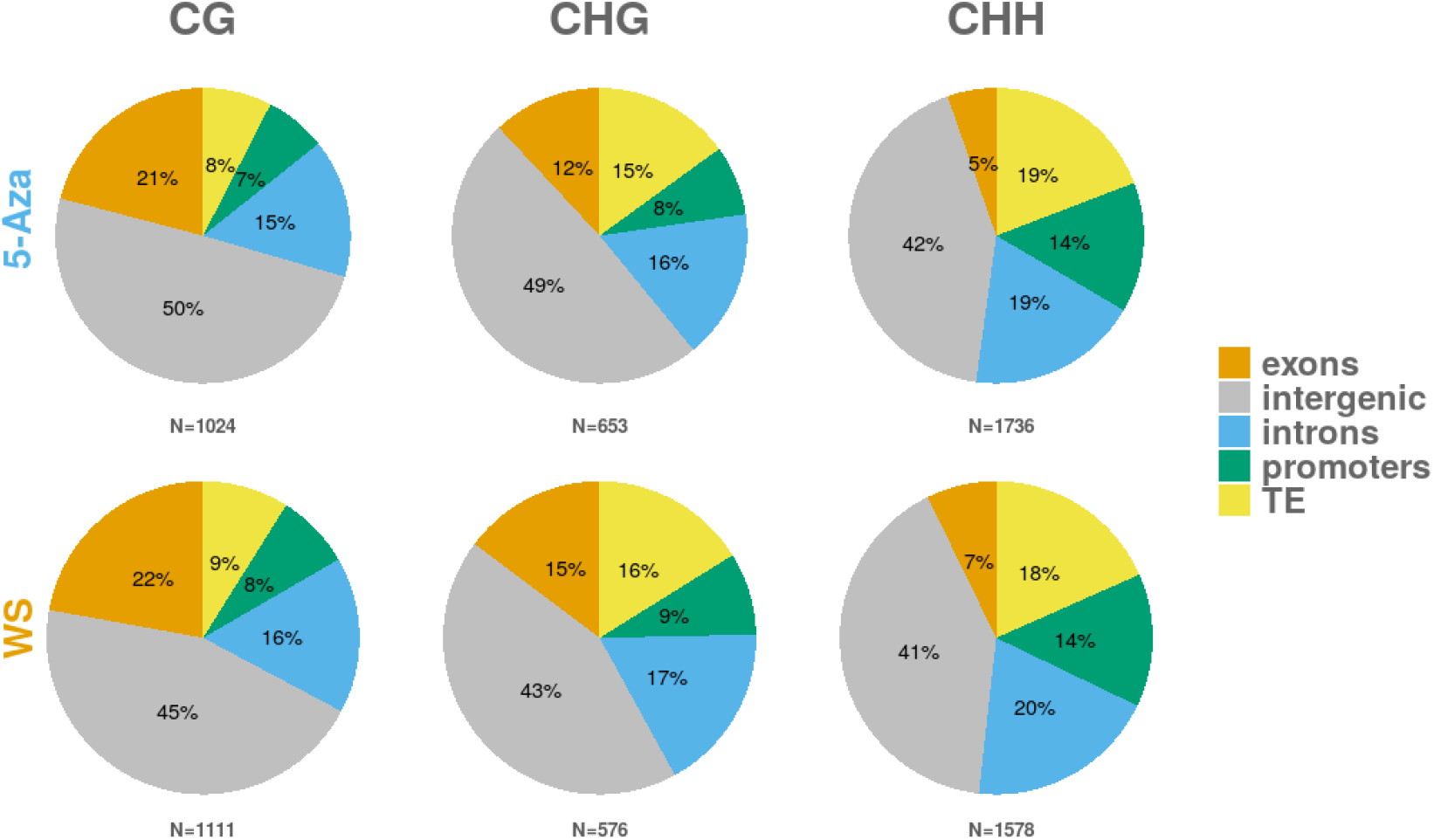
Distribution of the DMCs identified in 5-Azacytidine treatment (top) and water stress treatment (bottom). Exons, introns, and promoters corresponds to DMCs overlapping with genes, and TE corresponds to DMCs overlapping with TEs. All other remaining DMCs are classified as intergenic

**Fig. 5.**
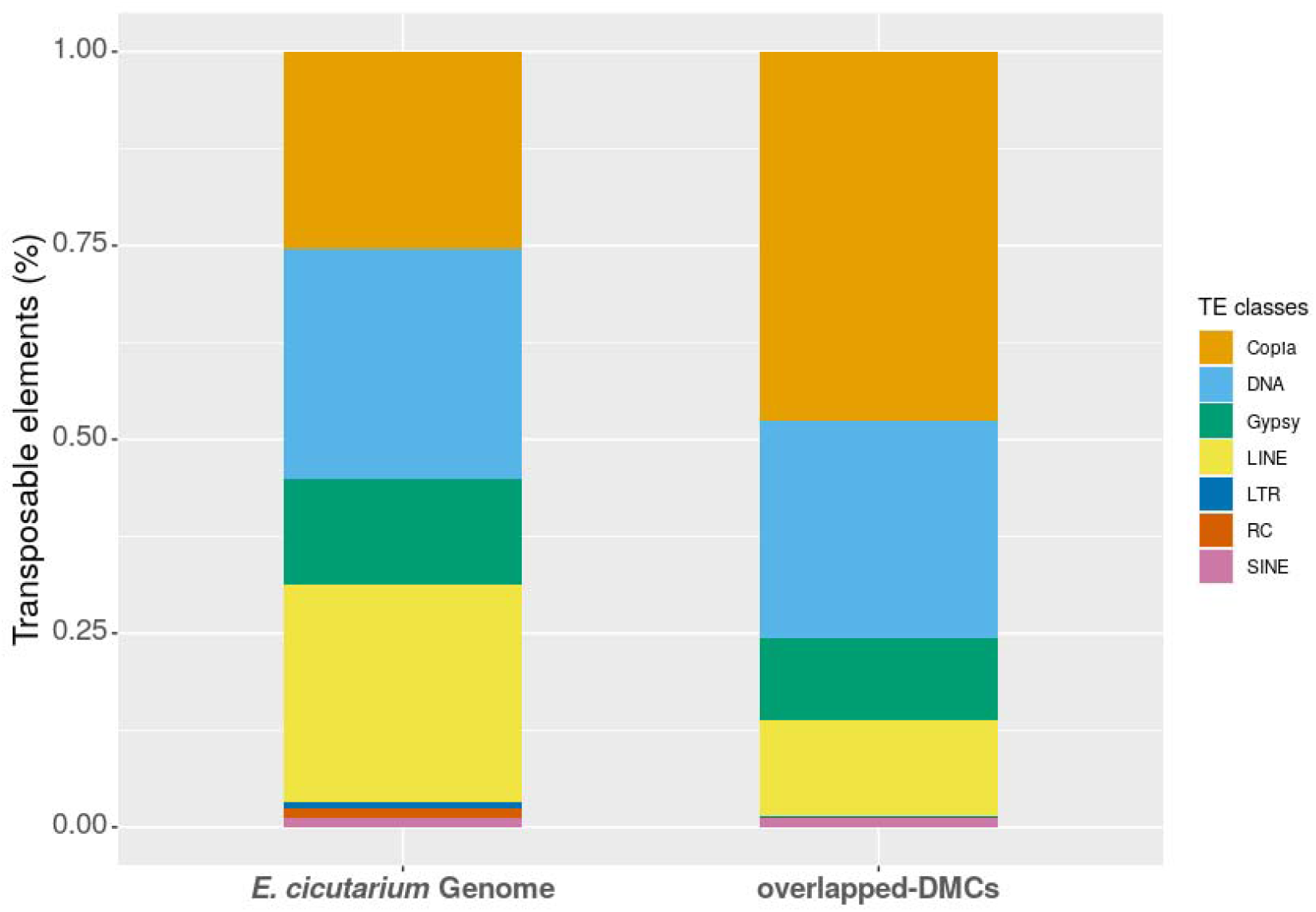
Distribution of the TE classes along *Erodium cicutarium* genome and distribution of TE classes for the overlapped-DMCs (N = 285,236 and 492, respectively).

Finally, although the disperse nature of the BsRADseq DMCs (Fig. S2) limits the possibility to identifying genomic regions with multiple DMCs (i.e., Differentially Methylation Regions, DMR), we obtained 29 potential DMRs covering gene regions (i.e., overlapping > 10 DMCs; Table 2). Eleven of these potential DMRs (37.9%) overlapped just intronic regions, whereas 31.0% appeared just on promotor regions and 13.8% just exonic regions. The annotation of these DMR-associated candidate genes included several with a potential role in stress response (Table 2).

**Table 2.** Annotation of gene regions overlapping DMRs (i.e., >= 10 DMCs). We showed the maximum number of DMCs, their genomic location, and the most frequent treatment (either 5-Aza or WS) or interaction cluster (denoted by C1-C8 as in Fig. 3) associated to DMCs obtained. See Table S2 for details.

| Gene | Annotation | DMCs |  |  | Treatment/<br>Cluster |
| --- | --- | --- | --- | --- | --- |
|  |  | Promoter | Exon | Intron |  |
| Ec-S10188_3 | CC-NBS-LRR resistance protein RGA3 | 28 | 0 | 0 | C1 |
| Ec-S25751_1 | Methyltransferase-like protein 13 | 19 | 0 | 0 | C5 |
| Ec-S20359_1 | ABC transporter F family member 4 | 17 | 0 | 0 | C3 |
| Ec-S27698_1 | Phenylalanine ammonia-lyase PAM | 17 | 0 | 0 | C6 |
| Ec-S71193_1 | Uncharacterized protein | 16 | 0 | 0 | C1 |
| Ec-S44030_4 | Cyclin-dependent protein kinases 2 CDKC | 13 | 0 | 0 | C1 |
| Ec-S06490_3 | Uncharacterized protein | 13 | 0 | 0 | C2 |
| Ec-S11263_1 | Bpg2-like protein (Fragment) | 11 | 0 | 0 | 5-Aza |
| Ec-S41184_1 | 1,4-alpha-D-glucan maltohydrolase | 10 | 0 | 1 | WS |
| Ec-S48520_2 | Uncharacterized protein | 6 | 6 | 0 | C1 |
| Ec-S09768_1 | Uncharacterized protein | 0 | 11 | 0 | C8 |
| Ec-S63962_1 | Reverse transcriptase (RNA-dependent DNA polymerase) | 0 | 10 | 0 | 5-Aza |
| Ec-S77906_2 | Uncharacterized protein | 0 | 10 | 0 | C1 |
| Ec-S53645_2 | Apoptosis inhibitory protein 5 (API5) | 0 | 10 | 0 | WS |
| Ec-S29620_1 | Amino acid permease 1 | 0 | 6 | 4 | WS |
| Ec-S77915_1 | ALE2 AtPERK1 | 0 | 1 | 18 | WS |
| Ec-S76299_2 | Agamous-like MADS-box protein AGL9 homolog | 0 | 1 | 17 | C6 |
| Ec-S71479_1 | Probable polygalacturonase | 0 | 1 | 9 | C3 |
| Ec-S06731_4 | DNA ligase 1 | 0 | 0 | 18 | C2 |
| Ec-S64428_1 | (1->3)-beta-glucan endohydrolase 10 | 0 | 0 | 16 | C1/C8 |
| Ec-S64428_1 | Glucan endo-1_3-beta-glucosidase 14 | 0 | 0 | 16 | C8 |
| Ec-S61275_5 | Uncharacterized protein | 0 | 0 | 14 | WS |
| Ec-S08686_1 | GDSL esterase/lipase At4g10955-like | 0 | 0 | 13 | WS |
| Ec-S63591_3 | Uncharacterized protein | 0 | 0 | 12 | C4 |
| Ec-S66538_1 | Uncharacterized protein | 0 | 0 | 12 | C5 |
| Ec-S76981_2 | hypothetical protein CCACVL1_13302 | 0 | 0 | 11 | C1 |
| Ec-S72001_2 | 60S ribosomal protein L37-3 | 0 | 0 | 10 | 5-Aza |
| Ec-S62651_1 | Uncharacterized protein | 0 | 0 | 10 | C4 |
| Ec-S28865_5 | Transcription factor WER-like | 0 | 0 | 10 | WS |

## Discussion

The broad interspecific variation in global DNA methylation level, its phylogenetic signal, and its positive relationship with the haploid genome size (Alonso et al., 2016; Niederhuth et al., 2016; Alonso et al., 2019), together with the evolution of plant specific methyltransferases (Bewick et al., 2017; Lyko, 2018; Kenchanmane Raju et al., 2019) support DNA methylation as a key component of plant epigenomes (Finnegan, 2010). DNA methylation status is regulated by *de novo* DNA methylation, maintenance of methylation at DNA replication, and active demethylation; these three processes together with the recruitment of histone modifiers and effectors contribute to genome stability during development, and in response to genomic and environmental stress (Law and Jacobsen, 2010; Boyko and Kovalchuk, 2011; Fitz-James and Cavalli, 2022). The deep molecular understanding of stress response in model plant species with well-annotated reference genomes contrasts the mostly indirect evidence gained for non-model plant species based on experimental alteration of DNA methylation and subsequent analysis of phenotypes, or from the analysis of methylation changes of anonymous markers, such as MSAP (Richards *et al*., 2017). Molecular insights gained by RRBS methods provide a bridge between these two approaches (Paun *et al*., 2019) and could inform whether epigenomic regulation in response to stress might act differently in species with contrasting epigenomic features relative to model organisms (Springer *et al*., 2016). In the following paragraphs we will discuss how our results can help understand the genomic signature of experimental demethylation conducted by application of 5-Aza at seed stage and its impact on plant methylome responses to recurrent water stress in an annual plant.

### Long-lasting methylation changes after seed demethylation detected by BsRADseq

5-Azacytidine is a structural analogue of 5-methyl-cytosine that can be incorporated into DNA where it establishes a covalent bond with a methyltransferase enzyme that can not be easily reversed. The enzyme is trapped reducing the number of active DNA methyltransferase enzymes in the cells and therefore passively inducing DNA demethylation (Griffin et al., 2016; Lopez et al., 2016; Lyko, 2018). Applied at seed stage, 5-Aza reduced global DNA methylation of seedlings in a dose-dependent manner (Griffin et al., 2016; Alonso et al., 2017). In *A. thaliana* seedlings the effect was more significant in CG context, where it reduced the frequency of fully methylated loci, particularly at highly methylated regions, whereas the loss of CHG and CHH methylation was mainly evident in highly methylated areas of the pericentromeric regions (Griffin et al., 2016).

Our analysis of changes in cytosine methylation in the DNA of *E. cicutarium* leaves collected at the onset of flowering is to the best of our knowledge the first attempt to characterize the long-lasting molecular signatures of seed demethylation treatment with 5-Aza. The abridged view of the methylome provided by BsRADseq analysis indicated a mild overall reduction in global cytosine methylation that was most evident in CHG contexts (Fig. 1). More interestingly, both hyper and hypoDMC were obtained in the leaf genomes of flowering plants treated with 5-Aza at seed stage in comparison to those grown from untreated seeds. In CG context the number of hypoDMCs was slightly higher, whereas the number of positions altered in either direction was similar at both CHG and CHH contexts. These findings altogether suggest that 5-Aza at seed stage significantly altered the patterning of cytosine methylation in genomes of adult plant, and might at least partially erase memories of past stress (see e.g., Akkerman *et al*., 2016). Seed demethylation is, thus, a suitable tool for analyzing the impact of deregulated epigenetic backgrounds on molecular and phenotypic response to stress analyzed either in the treated individuals or their progenies (Burton & Metcalfe, 2014). It is also important to highlight here the significant family effect detected by the analyses of global methylation level which reveals that experimental demethylation does not necessarily impact equally all treated individuals, an aspect that has been already reported in replicated ecological studies analyzing phenotypic effects of seed demethylation with different genetic lines or provenances (Herman *et al*., 2016; Troyee *et al*., 2022) in contrast to a single accession or genetic line (e.g., Akkerman *et al*., 2016). Thus, a homogeneous impact of the treatment should not be taken as granted and a screening of methylation changes across different genotypes, populations and species would be highly desirable to assess the actual impact of the 5-Aza seed treatment.

### Genomic methylation changes in response to recurrent water stress across the individual life cycle

Epigenetic response to abiotic stress can involve complex regulation of histone marks, small RNAs production and DNA methylation (Banerjee & Roychoudhury, 2017), that impinge transcription of metabolic pathways and, in some cases, can be retained as memory marks of past events to improve individual (Fleta-Soriano & Munné-Bosch, 2016) or transgenerational responses (Fitz-James & Cavalli, 2022). In particular, significant DNA methylation changes in response to moderate to heavy water stress treatments have been reported for several plant species (see e.g., Pardo *et al*., 2020 for several grasses; Sow *et al*., 2021 for poplar), including variable responses of crop varieties expected to have contrasting drought tolerance in *Vicia faba* (Abid *et al*., 2017) and *Oryza sativa* (Wang *et al*., 2016), supporting its relevance as a mechanism for both long-term wild plant adaptation and crop improvement. Mild water stress applied at early seedling stage has been found to alter DNA methylation in the leaves of *A. thaliana* Col-0 accession, mainly at CHH contexts and predominantly within TE sequences, and significantly up- and down-regulate transcription of multiple genes (Van Dooren *et al*., 2020). Some stress-induced epigenetic variants are reversible and recurrent stress is expected to promote molecular memory and increased tolerance in subsequent events (Wang *et al*., 2021). Our recurrent water stress, aimed to mimic unpredictable rain in Mediterranean ecosystems, led to a significant overall increased DNA cytosine methylation in all three contexts in genomes of *E. cicutarium* leaves at adult stage. Our analysis of DMCs indicated that a good fraction of cytosines across the genomes of plants experiencing WS were either hypo or hyper-methylated compared to those grown well-watered. A large proportion of DMCs in all contexts overlapped in our study intergenic regions (Fig. 4), an aspect in part potentially related to the reduced resolution of the annotation of the newly reported reference genome. Apart from that, methylation changes at CG contexts mainly overlapped exons, and those in CHH were found to be comparatively more frequently associated to TEs and introns.

### Combined effects of 5-Azacytidine and water stress on leaf methylome

Significant interactions between DNA demethylating agents and experimental stress treatments have been proposed as a reliable approach to investigate the role of DNA methylation in regulating plant responses to stress in non-model plant species (Puy *et al*., 2018; Alonso *et al*., 2019a). Phenotypic evidence of significant interaction effects in response to water stress has been gained in a few non-model plant species (Herman *et al*., 2016; Rendina González *et al*., 2016). Our study confirms a significant interaction at molecular level in the leaf genomes of *E. cicutarium* adult plants between 5-Aza-induced demethylation and water stress. This interaction appears to produce a complex epigenomic instability each factor making instable changes at the genome-wide scale. On one hand, 5-Aza application in seeds triggers widespread changes in DNA methylation in adult leaves, but importantly it likely alters the expression of transcriptional and post-transcriptional regulatory proteins as suggested by the DMRs overlapping the orthologous of *Methyltransferase 13, RNA-dependent DNA polymerase* and *60S ribosomal protein* (Litholdo & Bousquet-Antonelli, 2019; Sáez-Vásquez & Delseny, 2019). Interestingly, 5-Aza promoted the hypermethylation of the Ec-S10188_3 gene, which contained 28 differentially methylated cytosines in the promoter region (Table 2) and encodes a putative disease resistance protein, with high identity to Resistance genes analogs 3 (RGA3) in *Theobroma cacao* (Sekhwal *et al*., 2015). Likewise, the promoter of Ec-S44030_4 gene, a *Cyclin-dependent protein kinase (CDKC*) orthologous, related to stress tolerance mechanisms (Kitsios & Doonan, 2011), was also hypermethylated. Although, an antagonist effect of water stress and 5-Aza exposure was found for these genes (Cluster 1), the 5-Aza-induced hypermethylation may resemble a priming state for a faster future responses to disease attacks and abiotic stress (Latzel *et al*., 2016; Alonso *et al*., 2018).

On the other hand, environmental stresses, such as cold and drought, can also affect the methylase and/or demethylase activity (Lucibelli *et al*., 2022) explaining the interaction with the demethylation agent 5-Azacytidine. For example, heat stress induced downregulation of demethylases in *Triticum aestivum* (Gahlaut *et al*., 2022) but were significantly upregulated in *Camellia sinensis* under cold and drought (Zhu *et al*., 2020). Demethylases were also significantly induced by cold in *Dendrobium officinale* where methylases were also affected by drought (Yu *et al*., 2021). In our study, water stress, as well as the 5-Aza treatment, independently triggered the promoter hypomethylation of the *Methyltransferase 13* which may activate or upregulate its expression. However, the combination of treatments restored the control methylation level of this cytosine methylase. In addition, the methylation stage of many abiotic response-genes was directly altered by the water stress or its combination with the 5-Aza treatment. For example, the promoter regions of the abiotic stress related genes *Phenylalanine ammonia-lyase* (PAL, Jeong *et al*., 2012) and *ABC transporter F family* (ABCF, Dahuja *et al*., 2021) were also affected. Most methylation changes were observed at CHH context potentially linked to a genome-wide decrease in TE activity repression, as previously reported for other stress studies (e.g., Van Dooren *et al*., 2020).

### Changes in methylation in Transposable Elements

In plants, various TEs are involved in environmental stress adaptation (Casacuberta & González, 2013; Sahu *et al*., 2013). We found a significant differential methylation of *Copia* transposable elements in *Erodium cicutarium* under 5-Aza and WS treatments. *Copia* superfamily of long terminal repeat (LTR) retrotransposons is known to be activated by biotic and abiotic stresses (Ito, 2022). ONSEN elements are highly expressed under heat stress in Brassicaceae (Pietzenuk *et al*., 2016), whereas other *Copia* elements are activated by several environmental stresses in different species (see Table 1 in Ito, 2022). Transposons activation can affect gene regulation on a genome-wide scale by acting as new cis-regulatory elements and conferring increased stress-responsiveness to nearby protein-coding genes, which can facilitate acclimation or even adaptation to stressful conditions (Horváth *et al*., 2017; Dubin *et al*., 2018).Transposon activation under environmental stress can have negative effects on fitness (Horváth *et al*., 2017), but increasing TE methylation, especially in genomic regions neighbouring highly expressed stress-induced genes, could prevent TE transcription via RNA polymerase II and hence avoid deleterious effects upon the plant (Secco *et al*., 2015).

## Conclusions

By using a 2×2 factorial design, combining recurrent water stress and early life demethylation with a reduced representation method of surveying genome-wide methylation levels in adult plants, we show a significant interaction between methylation and plant stress response in the Mediterranean *Erodium cicutarium*. We uncover extensive effects of both seedling demethylation and water stress on adult DNA methylation, with a large portion of changes observed around genic regions and within TEs. Altogether, our results provide useful information towards understanding the role of DNA methylation in plant drought responses.

## Acknowledgements

We thank Laura Cabral, Esmeralda López, Daisy Johnson, Pablo Martín, Alejandro Mira, Noelia Zarza, and Elena Villa for assistance in greenhouse and laboratory; Maite Lorenzo for her training with BsRADseq wet lab; Rocío Esteban and all the staff of Allgenetics for their assistance with draft genome analyses; and Rubén Martín-Blázquez for suggestions on a draft version of the manuscript. We are particularly indebted to Carlos M. Herrera for his continuous encouragement and insightful discussions at all phases of this work. Financial support was provided by grants EPIECOL-CGL2013-43352-P, EPIENDEM-CGL2016-76605-P and EPINTER-PID2019-104365GB-I00 (Ministerio de Ciencia e Innovación, Spanish Government).

## Author contributions

C.A., F.B. and M.M. designed the research; C.A. and M.M. conducted greenhouse experiments; C.A., F.B. and O.P. designed the BsRADseq experiment and contributed to data analyses; P.B. prepared the libraries; F.B. led data analyses; C.A. led the writing; all authors contributed to refining the manuscript.

## Data availability

Raw data can be accessed in the Sequence Read Archive (SRA) with BioProject ID XXXXXXX.

## References

Abid G, Mingeot D, Muhovski Y, Mergeai G, Aouida M, Abdelkarim S, Aroua I, El Ayed M, M’hamdi M, Sassi K, et al. 2017. Analysis of DNA methylation patterns associated with drought stress response in faba bean (*Vicia faba* L.) using methylation-sensitive amplification polymorphism (MSAP). Environmental and Experimental Botany 142: 34–44.

Akkerman KC, Sattarin A, Kelly JK, Scoville AG. 2016. Transgenerational plasticity is sex-dependent and persistent in yellow monkeyflower (*Mimulus guttatus*). Environmental Epigenetics 2: 1–8.

Alioto T, Blanco E, Parra G, Guigó R. 2018. Using geneid to identify genes. Current Protocols in Bioinformatics 64: 1–32.

Alonso C, Medrano M, Pérez R, Bazaga P, Herrera CM. 2017. Tissue-specific response to experimental demethylation at seed germination in the non-model herb *Erodium cicutarium*. Epigenomes 1: 1–11.

Alonso C, Medrano M, Pérez R, Canto A, Parra-Tabla V, Herrera CM. 2019a. Interspecific variation across angiosperms in global DNA methylation: phylogeny, ecology and plant features in tropical and Mediterranean communities. New Phytologist 224:949–960.

Alonso C, Pérez R, Bazaga P, Medrano M, Herrera CM. 2016. MSAP markers and global cytosine methylation in plants: A literature survey and comparative analysis for a wild-growing species. Molecular Ecology Resources 16: 80–90.

Alonso C, Ramos-Cruz D, Becker C. 2018. The role of plant epigenetics in biotic interactions. New Phytologist.

Alonso C, Ramos-Cruz D, Becker C. 2019b. The role of plant epigenetics in biotic interactions. New Phytologist 221: 731–737.

Andrews KR, Good JM, Miller MR, Luikart G, Hohenlohe PA. 2016. Harnessing the power of RADseq for ecological and evolutionary genomics. Nature Reviews Genetics 17: 81–92.

Balao F, Paun O, Alonso C. 2018. Uncovering the contribution of epigenetics to plant phenotypic variation in Mediterranean ecosystems. Plant Biology 20: 38–49.

Banerjee A, Roychoudhury A. 2017. Epigenetic regulation during salinity and drought stress in plants: Histone modifications and DNA methylation. Plant Gene 11: 199–204.

Bäurle I. 2018. Can’t remember to forget you: Chromatin-based priming of somatic stress responses. Seminars in Cell and Developmental Biology 83: 133–139.

Burton T, Metcalfe NB. 2014. Can environmental conditions experienced in early life influence future generations? Proceedings of the Royal Society B: Biological Sciences 281.

Camacho C, Coulouris G, Avagyan V, Ma N, Papadopoulos J, Bealer K, Madden TL. 2009. BLAST+: Architecture and applications. BMC Bioinformatics 10: 1–9.

Casacuberta E, González J. 2013. The impact of transposable elements in environmental adaptation. Molecular Ecology 22: 1503–1517.

Chikhi R, Medvedev P. 2014. Informed and automated k-mer size selection for genome assembly. Bioinformatics 30: 31–37.

Cook BI, Anchukaitis KJ, Touchan R, Meko DM, Cook ER. 2016. Spatiotemporal drought variability in the mediterranean over the last 900 years. Journal of Geophysical Research 121:2060–2074.

Cowling RM, Ojeda F, Lamont BB, Rundel PW, Lechmere-Oertel R. 2005. Rainfall reliability, a neglected factor in explaining convergence and divergence of plant traits in fire-prone mediterranean-climate ecosystems. Global Ecology and Biogeography 14: 509–519.

Crisp PA, Ganguly D, Eichten SR, Borevitz JO, Pogson BJ. 2016. Reconsidering plant memory: Intersections between stress recovery, RNA turnover, and epigenetics. Science Advances 2.

Dahuja A, Kumar RR, Sakhare A, Watts A, Singh B, Goswami S, Sachdev A, Praveen S. 2021. Role of ATP-binding cassette transporters in maintaining plant homeostasis under abiotic and biotic stresses. Physiologia Plantarum 171: 785–801.

Dangi AK, Sharma B, Khangwal I, Shukla P. 2018. Combinatorial interactions of biotic and abiotic stresses in plants and their molecular mechanisms: systems biology approach. Molecular Biotechnology 60: 636–650.

Van Dooren TJM, Silveira AB, Gilbault E, Jiménez-Gómez JM, Martin A, Bach L, Tisné S, Quadrana L, Loudet O, Colot V. 2020. Mild drought in the vegetative stage induces phenotypic, gene expression, and DNA methylation plasticity in *Arabidopsis* but no transgenerational effects. Journal of Experimental Botany 71: 3588–3602.

Douma JC, Vermeulen PJ, Poelman EH, Dicke M, Anten NPR. 2017. When does it pay off to prime for defense? A modeling analysis. New Phytologist 216: 782–797.

Dubin MJ, Mittelsten Scheid O, Becker C. 2018. Transposons: a blessing curse. Current Opinion in Plant Biology 42: 23–29.

Feng H, Conneely KN, Wu H. 2014. A Bayesian hierarchical model to detect differentially methylated loci from single nucleotide resolution sequencing data. Nucleic Acids Research 42: 1–11.

Fitz-James MH, Cavalli G. 2022. Molecular mechanisms of transgenerational epigenetic inheritance. Nature Reviews Genetics 23: 325–341.

Fiz-Palacios O, Vargas P, Vila R, Papadopulos AST, Aldasoro JJ. 2010. The uneven phylogeny and biogeography of *Erodium* (Geraniaceae): Radiations in the Mediterranean and recent recurrent intercontinental colonization. Annals of Botany 106: 871–884.

Fleta-Soriano E, Munné-Bosch S. 2016. Stress memory and the inevitable effects of drought: A physiological perspective. Frontiers in Plant Science 7: 143.

Francis A, Darbyshire SJ, Légère A, Simard MJ. 2012. The biology of Canadian weeds. 151. *Erodium cicutarium* (L.) L’Hér. ex Aiton. Canadian Journal of Plant Science 92: 1359–1380.

Gahlaut V, Samtani H, Gautam T, Khurana P. 2022. Identification and characterization of DNA demethylase genes and their association with thermal stress in Wheat (*Triticum aestivum* L.). Frontiers in Genetics 13: 894020.

Gallusci P, Dai Z, Génard M, Gauffretau A, Leblanc-Fournier N, Richard-Molard C, Vile D, Brunel-Muguet S. 2017. Epigenetics for plant improvement: current knowledge and modeling avenues. Trends in Plant Science 22: 610–623.

Giorgi F, Lionello P. 2008. Climate change projections for the Mediterranean region. Global and Planetary Change 63: 90–104.

Griffin PT, Niederhuth CE, Schmitz RJ. 2016. A comparative analysis of 5-azacytidine-and zebularine-induced DNA demethylation. G3: Genes, Genomes, Genetics 6: 2773–2780.

Gutzat R, Mittelsten Scheid O. 2012. Epigenetic responses to stress: Triple defense? Current Opinion in Plant Biology 15: 568–573.

Herman JJ, Sultan SE, Herman JJ. 2016. DNA methylation mediates genetic variation for adaptive transgenerational plasticity. Proceedings of the Royal Society B: Biological Sciences 283: 20160988.

Herrera CM, Medrano M, Pérez R, Bazaga P, Alonso C. 2019. Within-plant heterogeneity in fecundity and herbivory induced by localized DNA hypomethylation in the perennial herb *Helleborus foetidus*. American Journal of Botany 106: 798–806.

Hirsch S, Baumberger R, Grossniklaus U. 2012. Epigenetic variation, inheritance, and selection in plant populations. Cold Spring Harbor Symposia on Quantitative Biology 77: 97–104.

Horváth V, Merenciano M, González J. 2017. Revisiting the Relationship between Transposable Elements and the Eukaryotic Stress Response. Trends in Genetics 33: 832–841.

IPCC. 2014. Climate Change 2014: Synthesis Report. Contribution of Working Groups I, II and III to the Fifth Assessment Report of the Intergovernmental Panel on Climate Change. Geneva, Switzerland.

Ito H. 2022. Environmental stress and transposons in plants. Genes & Genetic Systems: 22-00045.

Jeong MJ, Choi BS, Bae DW, Shin SC, Park SU, Lim HS, Kim J, Kim JB, Cho BK, Bae H. 2012. Differential expression of kenaf phenylalanine ammonia-lyase (PAL) ortholog during developmental stages and in response to abiotic stresses. Plant OMICS 5: 392–399.

Kitsios G, Doonan JH. 2011. Cyclin dependent protein kinases and stress responses in plants. Plant Signaling and Behavior 6: 204–209.

Kolde R. 2019. pheatmap: Pretty Heatmapsitle.: R package version 1.0.12.

Krueger F, Andrews SR. 2011. Bismark: A flexible aligner and methylation caller for Bisulfite-Seq applications. Bioinformatics 27: 1571–1572.

Langmead B, Salzberg SL. 2012. Fast gapped-read alignment with Bowtie 2. Nature methods 9: 357–9.

Latzel V. 2015. Pitfalls in ecological research – transgenerational effects. Folia Geobotanica 50: 75–85.

Latzel V, Rendina González AP, Rosenthal J. 2016. Epigenetic memory as a basis for intelligent behavior in clonal plants. Frontiers in Plant Science 7: 1–7.

Li H, Durbin R. 2010. Fast and accurate long-read alignment with Burrows-Wheeler transform. Bioinformatics 26: 589–595.

Li H, Handsaker B, Wysoker A, Fennell T, Ruan J, Homer N, Marth G, Abecasis G, Durbin R. 2009. The Sequence Alignment/Map format and SAMtools. Bioinformatics 25: 2078–2079.

Litholdo CG, Bousquet-Antonelli C. 2019. Chemical RNA modifications: The plant epitranscriptome. In: Epigenetics in Plants of Agronomic Importance: Fundamentals and Applications. Springer, 291–310.

López-Jurado J, Balao F, Mateos-Naranjo E. 2016. Deciphering the ecophysiological traits involved during water stress acclimation and recovery of the threatened wild carnation, Dianthus inoxianus. Plant Physiology and Biochemistry 109: 397–405.

López Rubio R, Pescador DS, Escudero A, Sánchez AM. 2022. Rainy years counteract negative effects of drought on taxonomic, functional, and phylogenetic diversity: resilience in annual plant communities. Journal of Ecology: 1–13.

Lopez M, Halby L, Arimondo PB. 2016. DNA methyltransferase inhibitors: Development and applications.

Lucibelli F, Valoroso MC, Aceto S. 2022. Plant DNA methylation □: an epigenetic mark in development, environmental interactions, and evolution. International Journal of Molecular Sciences 23: 8299.

Lyko F. 2018. The DNA methyltransferase family: A versatile toolkit for epigenetic regulation. Nature Reviews Genetics 19: 81–92.

Martin GT, Seymour DK, Gaut BS. 2021. CHH methylation islands: a nonconserved feature of grass genomes that is positively associated with transposable elements but negatively associated with gene-body methylation. Genome Biology and Evolution 13: 1–16.

Matesanz S, Valladares F. 2014. Ecological and evolutionary responses of Mediterranean plants to global change. Environmental and Experimental Botany 103: 53–67.

Mirouze M, Paszkowski J. 2011. Epigenetic contribution to stress adaptation in plants. Current Opinion in Plant Biology 14: 267–274.

Muñoz-Mérida A, Viguera E, Claros MG, Trelles O, Pérez-Pulido AJ. 2014. Sma3s: A three-step modular annotator for large sequence datasets. DNA Research 21: 341–353.

Niederhuth CE, Schmitz RJ. 2017. Putting DNA methylation in context: from genomes to gene expression in plants. Biochimica et Biophysica Acta - Gene Regulatory Mechanisms 1860: 149–156.

Pandey P, Ramegowda V, Senthil-Kumar M. 2015. Shared and unique responses of plants to multiple individual stresses and stress combinations: Physiological and molecular mechanisms. Frontiers in Plant Science 6: 1–14.

Pardo J, Wai CM, Chay H, Madden CF, Hilhorst HWM, Farrant JM, VanBuren R. 2020. Intertwined signatures of desiccation and drought tolerance in grasses. Proceedings of the National Academy of Sciences of the United States of America 117: 10079–10088.

Paun O, Verhoeven KJF, Richards CL. 2019. Opportunities and limitations of reduced representation bisulfite sequencing in plant ecological epigenomics. New Phytologist 221: 738–742.

Peng H, Zhang J. 2009. Plant genomic DNA methylation in response to stresses: Potential applications and challenges in plant breeding. Progress in Natural Science 19: 1037–1045.

Pietzenuk B, Markus C, Gaubert H, Bagwan N, Merotto A, Bucher E, Pecinka A. 2016. Recurrent evolution of heat-responsiveness in Brassicaceae COPIA elements. Genome Biology 17: 1–15.

Pustahija F, Brown SC, Bogunić F, Bašić N, Muratović E, Ollier S, Hidalgo O, Bourge M, Stevanovic V, Siljak-Yakovlev S. 2013. Small genomes dominate in plants growing on serpentine soils in West Balkans, an exhaustive study of 8 habitats covering 308 taxa. Plant and Soil 373: 427–453.

Puy J, Dvořáková H, Carmona CP, de Bello F, Hiiesalu I, Latzel V. 2018. Improved demethylation in ecological epigenetic experiments: Testing a simple and harmless foliar demethylation application. Methods in Ecology and Evolution 9: 744–753.

R Core Team. 2022. A language and environment for statistical computing.

Renaud G, Stenzel U, Maricic T, Wiebe V, Kelso J. 2015. DeML: Robust demultiplexing of Illumina sequences using a likelihood-based approach. Bioinformatics 31: 770–772.

Rendina González AP, Chrtek J, Dobrev PI, Dumalasová V, Fehrer J, Mráz P, Latzel V. 2016. Stress-induced memory alters growth of clonal offspring of white clover (*Trifolium repens*). American Journal of Botany 103: 1567–1574.

Richards CL, Alonso C, Becker C, Bossdorf O, Bucher E, Colomé-Tatché M, Durka W, Engelhardt J, Gaspar B, Gogol-Döring A, et al. 2017. Ecological plant epigenetics: Evidence from model and non-model species, and the way forward. Ecology Letters 20: 1576–1590.

Rochette NC, Rivera-Colón AG, Catchen JM. 2019. Stacks 2: Analytical methods for paired-end sequencing improve RADseq-based population genomics. Molecular Ecology 28: 4737–4754.

Sáez-Vásquez J, Delseny M. 2019. Ribosome biogenesis in plants: From functional 45S ribosomal DNA organization to ribosome assembly factors. Plant Cell 31: 1945–1967.

Sahu PP, Pandey G, Sharma N, Puranik S, Muthamilarasan M, Prasad M. 2013. Epigenetic mechanisms of plant stress responses and adaptation. Plant Cell Reports 32: 1151–1159.

Schmidt M, Van Bel M, Woloszynska M, Slabbinck B, Martens C, De Block M, Coppens F, Van Lijsebettens M. 2017. Plant-RRBS, a bisulfite and next-generation sequencing-based methylome profiling method enriching for coverage of cytosine positions. BMC Plant Biology 17: 1–14.

Secco D, Wang C, Shou H, Schultz MD, Chiarenza S, Nussaume L, Ecker JR, Whelan J, Lister R. 2015. Stress induced gene expression drives transient DNA methylation changes at adjacent repetitive elements. eLife 4: 1–26.

Sekhwal MK, Li P, Lam I, Wang X, Cloutier S, You FM. 2015. Disease resistance gene analogs (RGAs) in plants. International Journal of Molecular Sciences 16: 19248–19290.

Simpson JT, Wong K, Jackman SD, Schein JE, Jones SJM, Birol I. 2009. ABySS: A parallel assembler for short read sequence data. Genome Research 19: 1117–1123.

Sow MD, Le Gac AL, Fichot R, Lanciano S, Delaunay A, Le Jan I, Lesage-Descauses MC, Citerne S, Caius J, Brunaud V, et al. 2021. RNAi suppression of DNA methylation affects the drought stress response and genome integrity in transgenic poplar. New Phytologist 232: 80–97.

Springer NM, Lisch D, Li Q. 2016. Creating order from chaos: epigenome dynamics in plants with complex genomes. The Plant Cell 28: 314–325.

Springer NM, Schmitz RJ. 2017. Exploiting induced and natural epigenetic variation for crop improvement. Nature Publishing Group.

Sun L, Jing Y, Liu X, Li Q, Xue Z, Cheng Z, Wang D, He H, Qian W. 2020. Heat stress-induced transposon activation correlates with 3D chromatin organization rearrangement in Arabidopsis. Nature Communications 11.

The Uniprot Consortium. 2007. The Universal Protein Resource (UniProt). Nucleic Acids Research 35: 193–197.

Tricker PJ. 2015. Transgenerational inheritance or resetting of stress-induced epigenetic modifications: Two sides of the same coin. Frontiers in Plant Science 6: 1–6.

Troyee AN, Medrano M, Müller C, Alonso C. 2022. Variation in DNA Methylation and Response to Short-Term Herbivory in *Thlaspi Arvense*. Flora 293: 152106.

Trucchi E, Mazzarella AB, Gilfillan GD, Lorenzo MT, Schönswetter P, Paun O. 2016. BsRADseq: screening DNA methylation in natural populations of non-model species. Molecular Ecology 25: 1697–1713.

Verhoeven KJF, Jansen JJ, van Dijk PJ, Biere A. 2010. Stress-induced DNA methylation changes and their heritability in asexual dandelions. New Phytologist 185: 1108–1118.

Verhoeven KJF, Preite V. 2014. Epigenetic variation in asexually reproducing organisms. Evolution 68: 644–655.

Walter J, Jentsch A, Beierkuhnlein C, Kreyling J. 2013. Ecological stress memory and cross stress tolerance in plants in the face of climate extremes. Environmental and Experimental Botany 94: 3–8.

Walter J, Nagy L, Hein R, Rascher U, Beierkuhnlein C, Willner E, Jentsch A. 2011. Do plants remember drought? Hints towards a drought-memory in grasses. Environmental and Experimental Botany 71: 34–40.

Wang L, Cao S, Wang P, Lu K, Song Q, Zhao FJ, Chen ZJ. 2021. DNA hypomethylation in tetraploid rice potentiates stress-responsive gene expression for salt tolerance. Proceedings of the National Academy of Sciences of the United States of America 118: 1–10.

Wang W, Qin Q, Sun F, Wang Y, Xu D, Li Z, Fu B. 2016. Genome-wide differences in DNA methylation changes in two contrasting rice genotypes in response to drought conditions. Frontiers in Plant Science 7: 1–13.

Waterhouse RM, Seppey M, Simao FA, Manni M, Ioannidis P, Klioutchnikov G, Kriventseva E V., Zdobnov EM. 2018. BUSCO applications from quality assessments to gene prediction and phylogenomics. Molecular Biology and Evolution 35: 543–548.

Witten DM, Tibshirani R. 2010. A framework for feature selection in clustering. Journal of the American Statistical Association 105: 713–726.

Yu Z, Zhang G, Teixeira da Silva JA, Li M, Zhao C, He C, Si C, Zhang M, Duan J. 2021. Genome-wide identification and analysis of DNA methyltransferase and demethylase gene families in *Dendrobium officinale* reveal their potential functions in polysaccharide accumulation. BMC Plant Biology 21: 1–17.

Zhang X, Zwiers FW, Hegerl GC, Lambert FH, Gillett NP, Solomon S, Stott PA, Nozawa T. 2007. Detection of human influence on twentieth-century precipitation trends. Nature 448: 461–465.

Zhu C, Zhang S, Zhou C, Chen L, Fu H, Li X, Lin Y, Lai Z, Guo Y. 2020. Genome-wide investigation and transcriptional analysis of cytosine-5 DNA methyltransferase and DNA demethylase gene families in tea plant (*Camellia sinensis*) under abiotic stress and withering processing. PeerJ 2020.

