## Supplementary materials for "Long-term methylome changes after experimental seed demethylation and their interaction with recurrent water stress in *Erodium cicutarium* (Geraniaceae)"

**Table S1**. Summary of sequencing and mapping results of the 32 individuals included in the bsRAD-seq library analysed.

| Treatment | | | | | | | | | |
|---|---|---|---|---|---|---|---|---|---|
| 5-Aza | Water | Family | Sample | No. clean paired-reads | Low quality (%) | GC content (%) | BS conversion rate (%) | Mapping rate (%) | Treatment combination level |
| C | WW | PT_1031 | CC_253 | 4932295 | 0.16 | 27.5 | 99.51 | 40.2 | Control |
| C | WW | PT_1001 | CC_225 | 5907482 | 0.16 | 28.5 | 99.33 | 39.7 | Control |
| C | WW | PT_0969 | CC_147 | 2775753 | 0.17 | 28.0 | 99.21 | 36.8 | Control |
| C | WW | PT_0925 | CC_001 | 7194273 | 0.16 | 29.5 | 99.43 | 36.9 | Control |
| C | WW | CH_1307 | CC_491 | 1976553 | 0.15 | 29.0 | 99.45 | 42.2 | Control |
| C | WW | CH_1245 | CC_433 | 1367138 | 0.15 | 29.5 | 99.56 | 35.1 | Control |
| C | WW | CH_1233 | CC_409 | 5295301 | 0.16 | 26.5 | 99.42 | 39.0 | Control |
| C | WW | CH_1187 | CC_291 | 5193693 | 0.17 | 30.5 | 99.46 | 35.2 | Control |
| C | WS | PT_1031 | CD_259 | 6767212 | 0.16 | 26.5 | 99.36 | 41.0 | WS |
| C | WS | PT_1001 | CD_219 | 6453453 | 0.16 | 31.0 | 99.31 | 37.6 | WS |
| C | WS | PT_0969 | CD_163 | 7364511 | 0.16 | 28.0 | 99.45 | 37.5 | WS |
| C | WS | PT_0925 | CD_005 | 11417213 | 0.16 | 28.0 | 99.40 | 39.5 | WS |
| C | WS | CH_1307 | CD_485 | 1133391 | 0.15 | 28.5 | 99.58 | 41.4 | WS |
| C | WS | CH_1245 | CD_445 | 1676382 | 0.15 | 28.0 | 99.27 | 38.0 | WS |
| C | WS | CH_1233 | CD_431 | 4738894 | 0.16 | 27.0 | 99.55 | 40.2 | WS |
| C | WS | CH_1187 | CD_297 | 3040251 | 0.17 | 31.0 | 99.40 | 35.0 | WS |
| A | WW | PT_1031 | AC_252 | 8958157 | 0.16 | 29.0 | 99.47 | 38.9 | 5-Aza |
| A | WW | PT_1001 | AC_220 | 5595964 | 0.17 | 28.0 | 99.40 | 37.7 | 5-Aza |
| A | WW | PT_0969 | AC_146 | 2702015 | 0.17 | 28.5 | 99.48 | 36.6 | 5-Aza |
| A | WW | PT_0925 | AC_016 | 4224209 | 0.16 | 31.0 | 99.23 | 37.3 | 5-Aza |
| A | WW | CH_1307 | AC_494 | 2236321 | 0.15 | 28.0 | 99.25 | 42.0 | 5-Aza |
| A | WW | CH_1245 | AC_438 | 1209208 | 0.15 | 30.0 | 99.47 | 37.5 | 5-Aza |
| A | WW | CH_1233 | AC_422 | 5638260 | 0.16 | 28.0 | 99.38 | 38.3 | 5-Aza |
| A | WW | CH_1187 | AC_300 | 3324116 | 0.16 | 28.0 | 99.56 | 36.3 | 5-Aza |
| A | WS | PT_1031 | AD_254 | 7102151 | 0.16 | 29.5 | 99.53 | 41.2 | 5-Aza+WS |
| A | WS | PT_1001 | AD_222 | 2275920 | 0.17 | 31.0 | 99.59 | 36.5 | 5-Aza+WS |
| A | WS | PT_0969 | AD_164 | 4611977 | 0.16 | 29.0 | 99.47 | 38.6 | 5-Aza+WS |
| A | WS | PT_0925 | AD_020 | 5908542 | 0.16 | 30.0 | 99.24 | 37.5 | 5-Aza+WS |
| A | WS | CH_1307 | AD_482 | 1931984 | 0.15 | 28.0 | 99.40 | 42.5 | 5-Aza+WS |
| A | WS | CH_1245 | AD_436 | 1926343 | 0.16 | 30.0 | 99.45 | 36.7 | 5-Aza+WS |
| A | WS | CH_1233 | AD_416 | 4371719 | 0.17 | 29.0 | 99.18 | 39.1 | 5-Aza+WS |
| A | WS | CH_1187 | AD_292 | 4316573 | 0.17 | 31.0 | 99.49 | 35.7 | 5-Aza+WS |

**Table S2**. Details of the DMR (≥ 10 DMC) in *Erodium cicutarium* (Average methylation by treatment, q-values and cluster)

| Loci | Context | Control | 5-Aza | WS | 5-AzaWS | Qval_5Aza | Qval_WS | Qval_5AzaWS | Treatment/ Cluster |
|---|---|---|---|---|---|---|---|---|---|
| *S10188_13237* | CHG | 0.000 | 1.000 | 0.000 | 0.000 | 1.26E-119 | 7.89E-102 | 1.19E-64 | C1 |
| *S10188_13239* | CHH | 0.000 | 1.000 | 0.000 | 0.000 | 3.24E-119 | 2.56E-101 | 3.58E-64 | C1 |
| *S10188_13240* | CHH | 0.000 | 1.000 | 0.000 | 0.000 | 3.24E-119 | 2.56E-101 | 3.58E-64 | C1 |
| *S10188_13241* | CHH | 0.000 | 1.000 | 0.000 | 0.000 | 3.24E-119 | 2.56E-101 | 3.58E-64 | C1 |
| *S10188_13243* | CHH | 0.000 | 1.000 | 0.000 | 0.000 | 3.24E-119 | 2.56E-101 | 3.58E-64 | C1 |
| *S10188_13246* | CG | 0.000 | 1.000 | 0.000 | 0.000 | 8.87E-120 | 7.29E-102 | 9.31E-65 | C1 |
| *S10188_13248* | CHH | 0.000 | 1.000 | 0.000 | 0.000 | 3.24E-119 | 2.56E-101 | 3.58E-64 | C1 |
| *S10188_13249* | CHH | 0.000 | 1.000 | 0.000 | 0.000 | 3.24E-119 | 2.56E-101 | 3.58E-64 | C1 |
| *S10188_13251* | CHH | 0.000 | 1.000 | 0.000 | 0.000 | 3.24E-119 | 2.56E-101 | 3.58E-64 | C1 |
| *S10188_13256* | CHH | 0.000 | 1.000 | 0.000 | 0.000 | 3.24E-119 | 2.56E-101 | 3.58E-64 | C1 |
| *S10188_13277* | CHH | 0.000 | 1.000 | 0.000 | 0.000 | 3.24E-119 | 2.56E-101 | 3.58E-64 | C1 |
| *S10188_13283* | CG | 0.000 | 1.000 | 0.000 | 0.000 | 1.10E-119 | 1.01E-101 | 1.15E-64 | C1 |
| *S10188_13286* | CHH | 0.000 | 1.000 | 0.000 | 0.000 | 3.24E-119 | 2.56E-101 | 3.58E-64 | C1 |
| *S10188_13287* | CHH | 0.000 | 1.000 | 0.000 | 0.000 | 3.24E-119 | 2.56E-101 | 3.58E-64 | C1 |
| *S10188_13288* | CHH | 0.000 | 1.000 | 0.000 | 0.000 | 3.24E-119 | 2.56E-101 | 3.58E-64 | C1 |
| *S10188_13297* | CG | 0.004 | 1.000 | 0.000 | 0.000 | 3.42E-116 | 7.29E-102 | 9.28E-63 | C1 |
| *S10188_13303* | CHH | 0.000 | 1.000 | 0.000 | 0.000 | 3.24E-119 | 2.56E-101 | 3.58E-64 | C1 |
| *S10188_13308* | CHH | 0.000 | 1.000 | 0.000 | 0.000 | 3.24E-119 | 2.56E-101 | 4.41E-64 | C1 |
| *S10188_13316* | CHH | 0.000 | 1.000 | 0.000 | 0.000 | 3.73E-119 | 2.56E-101 | 3.75E-64 | C1 |
| *S10188_13318* | CHH | 0.000 | 1.000 | 0.000 | 0.000 | 3.24E-119 | 2.56E-101 | 3.58E-64 | C1 |
| *S10188_13327* | CG | 0.000 | 0.963 | 0.007 | 0.000 | 4.50E-94 | 4.35E-80 | 1.41E-55 | C1 |
| *S10188_13336* | CHH | 0.000 | 1.000 | 0.000 | 0.000 | 3.24E-119 | 2.56E-101 | 3.58E-64 | C1 |
| *S10188_13339* | CHH | 0.000 | 1.000 | 0.000 | 0.000 | 3.24E-119 | 2.56E-101 | 3.58E-64 | C1 |
| *S10188_13351* | CHH | 0.000 | 1.000 | 0.000 | 0.000 | 4.29E-119 | 2.56E-101 | 3.92E-64 | C1 |
| *S10188_13357* | CHH | 0.000 | 1.000 | 0.000 | 0.000 | 3.24E-119 | 2.56E-101 | 3.58E-64 | C1 |
| *S10188_13362* | CG | 0.000 | 1.000 | 0.000 | 0.000 | 6.70E-120 | 5.29E-102 | 7.42E-65 | C1 |
| *S10188_13365* | CHH | 0.000 | 1.000 | 0.000 | 0.000 | 3.64E-119 | 2.56E-101 | 3.72E-64 | C1 |
| *S10188_13368* | CHH | 0.000 | 1.000 | 0.000 | 0.000 | 3.24E-119 | 2.56E-101 | 3.58E-64 | C1 |

| | | | | | | | | | |
|---|---|---|---|---|---|---|---|---|---|
| *S11263_305* | CHH | 1.000 | 0.500 | 0.748 | 1.000 | 1.97E-10 | 1.00E+00 | 1.00E+00 | 5-Aza |
| *S11263_312* | CG | 1.000 | 0.500 | 1.000 | 1.000 | 8.58E-24 | 1.00E+00 | 1.00E+00 | 5-Aza |
| *S11263_321* | CHG | 0.980 | 0.500 | 0.745 | 1.000 | 7.93E-10 | 1.00E+00 | 1.00E+00 | 5-Aza |
| *S11263_324* | CHG | 1.000 | 0.500 | 0.750 | 1.000 | 9.88E-12 | 1.00E+00 | 1.00E+00 | 5-Aza |
| *S11263_328* | CHG | 1.000 | 0.500 | 0.750 | 1.000 | 9.88E-12 | 1.00E+00 | 1.00E+00 | 5-Aza |
| *S11263_333* | CHG | 1.000 | 0.500 | 0.750 | 1.000 | 9.88E-12 | 1.00E+00 | 1.00E+00 | 5-Aza |
| *S11263_340* | CHG | 1.000 | 0.500 | 0.750 | 1.000 | 9.88E-12 | 1.00E+00 | 1.00E+00 | 5-Aza |
| *S11263_343* | CHH | 1.000 | 0.500 | 0.000 | 0.000 | 4.20E-23 | 1.00E+00 | 1.00E+00 | 5-Aza |
| *S11263_353* | CHG | 1.000 | 0.500 | 0.750 | 1.000 | 9.88E-12 | 1.00E+00 | 1.00E+00 | 5-Aza |
| *S11263_381* | CHH | 1.000 | 0.000 | 0.000 | 0.000 | 1.68E-32 | 1.00E+00 | 1.00E+00 | 5-Aza |
| *S11263_384* | CHH | 1.000 | 0.000 | 0.000 | 0.000 | 1.68E-32 | 1.00E+00 | 1.00E+00 | 5-Aza |
| *S20359_1400* | CG | 0.333 | 0.750 | 0.000 | 0.333 | 9.22E-15 | 1.48E-15 | 2.95E-02 | C1 |
| *S20359_1404* | CG | 0.333 | 0.750 | 0.000 | 0.333 | 9.22E-15 | 1.48E-15 | 2.95E-02 | C1 |
| *S20359_1434* | CG | 0.333 | 0.749 | 0.333 | 0.222 | 1.01E-15 | 3.07E-18 | 8.22E-04 | C1 |
| *S20359_1446* | CG | 0.999 | 0.999 | 0.562 | 0.667 | 1.00E+00 | 1.00E+00 | 6.89E-22 | C7 |
| *S20359_1463* | CHH | 0.323 | 0.250 | 0.000 | 0.330 | 1.59E-02 | 8.65E-40 | 1.39E-36 | C3 |
| *S20359_1464* | CHH | 0.216 | 0.250 | 0.000 | 0.333 | 2.48E-02 | 5.22E-44 | 3.02E-39 | C3 |
| *S20359_1465* | CHH | 0.000 | 0.250 | 0.000 | 0.333 | 1.00E+00 | 3.00E-56 | 3.08E-28 | C3 |
| *S20359_1491* | CHH | 0.000 | 0.250 | 0.562 | 0.333 | 1.00E+00 | 7.45E-44 | 4.47E-01 | WS |
| *S20359_1493* | CHH | 1.000 | 0.000 | 0.464 | 0.167 | 4.25E-109 | 1.00E+00 | 8.46E-10 | C4 |
| *S20359_1504* | CHG | 1.000 | 1.000 | 0.797 | 0.833 | 1.00E+00 | 1.00E+00 | 1.11E-08 | C7 |
| *S20359_1522* | CHH | 0.750 | 0.250 | 0.797 | 0.500 | 1.56E-38 | 1.74E-41 | 2.06E-34 | C5 |
| *S20359_1530* | CG | 1.000 | 1.000 | 0.464 | 875.000 | 1.00E+00 | 1.00E+00 | 3.82E-13 | C6 |
| *S20359_1532* | CHH | 0.750 | 375.000 | 0.698 | 625.000 | 1.97E-37 | 9.20E-71 | 1.48E-43 | C5 |
| *S20359_1533* | CHH | 0.750 | 375.000 | 0.696 | 625.000 | 2.34E-37 | 1.37E-70 | 1.27E-43 | C5 |
| *S20359_1548* | CHH | 0.250 | 375.000 | 0.001 | 0.657 | 1.00E+00 | 3.77E-182 | 1.71E-58 | C3 |
| *S20359_1553* | CHH | 0.000 | 375.000 | 0.001 | 0.667 | 1.00E+00 | 1.39E-297 | 7.54E-84 | C3 |
| *S20359_1554* | CHH | 0.000 | 375.000 | 0.000 | 0.665 | 1.00E+00 | 6.92E-287 | 1.99E-140 | C3 |
| *S25751_3063* | CG | 1.000 | 0.000 | 1.000 | 1.000 | 3.86E-18 | 1.17E-12 | 9.40E-10 | C5 |
| *S25751_3066* | CHH | 0.000 | 0.000 | 1.000 | 0.500 | 1.00E+00 | 2.45E-11 | 1.00E+00 | WS |
| *S25751_3073* | CG | 0.500 | 0.000 | 1.000 | 1.000 | 2.29E-13 | 9.90E-11 | 1.71E-06 | C5 |

| | | | | | | | | | |
|---|---|---|---|---|---|---|---|---|---|
| *S25751_3084* | CHH | 0.998 | 0.000 | 0.000 | 1.000 | 3.39E-16 | 5.71E-12 | 4.64E-46 | C6 |
| *S25751_3091* | CHH | 0.500 | 0.000 | 0.000 | 0.500 | 1.00E+00 | 1.98E-06 | 3.36E-04 | C3 |
| *S25751_3093* | CHH | 1.000 | 0.000 | 0.000 | 1.000 | 1.95E-17 | 5.71E-12 | 1.42E-47 | C6 |
| *S25751_3101* | CG | 0.998 | 0.000 | 1.000 | 1.000 | 7.01E-17 | 1.17E-12 | 5.48E-09 | C5 |
| *S25751_3104* | CG | 0.500 | 0.000 | 1.000 | 0.500 | 1.00E+00 | 4.03E-07 | 1.00E+00 | WS |
| *S25751_3107* | CHH | 1.000 | 0.000 | 1.000 | 1.000 | 1.89E-17 | 5.71E-12 | 4.61E-09 | C5 |
| *S25751_3114* | CHH | 1.000 | 0.000 | 1.000 | 1.000 | 1.89E-17 | 5.71E-12 | 4.61E-09 | C5 |
| *S25751_3122* | CHH | 0.000 | 0.000 | 1.000 | 0.500 | 1.00E+00 | 2.45E-11 | 1.00E+00 | WS |
| *S25751_3127* | CHG | 1.000 | 0.000 | 0.000 | 0.500 | 5.82E-18 | 1.00E+00 | 1.90E-10 | C4 |
| *S25751_3131* | CHG | 1.000 | 0.000 | 1.000 | 1.000 | 5.82E-18 | 1.76E-12 | 1.42E-09 | C5 |
| *S25751_3136* | CHH | 1.000 | 1.000 | 1.000 | 0.500 | 1.00E+00 | 2.45E-11 | 1.46E-07 | C7 |
| *S25751_3142* | CHH | 0.500 | 0.000 | 1.000 | 0.500 | 1.58E-07 | 1.00E+00 | 1.00E+00 | 5-Aza |
| *S25751_3144* | CG | 1.000 | 0.000 | 1.000 | 1.000 | 3.86E-18 | 1.17E-12 | 9.40E-10 | C5 |
| *S25751_3146* | CHH | 0.500 | 0.000 | 1.000 | 0.500 | 1.58E-07 | 1.00E+00 | 1.00E+00 | 5-Aza |
| *S25751_3149* | CG | 1.000 | 1.000 | 0.000 | 1.000 | 1.00E+00 | 1.00E+00 | 3.57E-11 | C6 |
| *S25751_3150* | CHG | 1.000 | 0.000 | 1.000 | 0.500 | 5.82E-18 | 1.00E+00 | 1.00E+00 | NA |
| *S27698_1141* | CHH | 0.000 | 1.000 | 0.000 | 1.000 | 2.48E-130 | 1.00E+00 | 1.00E+00 | NA |
| *S27698_1148* | CHG | 0.438 | 1.000 | 0.000 | 1.000 | 8.39E-04 | 1.00E+00 | 9.95E-49 | C6 |
| *S27698_1172* | CHH | 0.056 | 0.265 | 0.000 | 0.000 | 7.31E-13 | 5.33E-64 | 8.56E-06 | C1 |
| *S27698_1174* | CHH | 0.993 | 1.000 | 0.000 | 1.000 | 1.00E+00 | 1.00E+00 | 2.52E-81 | C6 |
| *S27698_1175* | CHH | 1.000 | 0.999 | 0.000 | 1.000 | 1.00E+00 | 1.00E+00 | 2.56E-97 | C6 |
| *S27698_1179* | CHH | 0.429 | 0.266 | 0.000 | 1.000 | 6.60E-06 | 2.67E-54 | 6.41E-145 | C3 |
| *S27698_1180* | CHH | 1.000 | 1.000 | 0.000 | 0.000 | 1.00E+00 | 9.62E-247 | 1.00E+00 | WS |
| *S27698_1183* | CHH | 0.559 | 0.737 | 0.000 | 1.000 | 3.33E-08 | 4.29E-63 | 6.08E-50 | C6 |
| *S27698_1185* | CHH | 0.438 | 0.264 | 0.000 | 1.000 | 1.35E-07 | 4.31E-55 | 7.89E-151 | C3 |
| *S27698_1189* | CHH | 1.000 | 0.262 | 0.000 | 1.000 | 7.12E-36 | 1.62E-55 | 1.23E-210 | C6 |
| *S27698_1236* | CHH | 0.438 | 0.736 | 1.000 | 1.000 | 1.12E-10 | 2.84E-63 | 1.62E-04 | C8 |
| *S27698_1238* | CHH | 0.438 | 0.997 | 1.000 | 1.000 | 1.13E-02 | 1.00E+00 | 1.00E+00 | 5-Aza |
| *S27698_1250* | CHH | 0.438 | 0.265 | 0.000 | 0.000 | 2.25E-07 | 5.33E-64 | 1.28E-02 | C4 |
| *S27698_1254* | CHG | 1.000 | 0.003 | 0.997 | 1.000 | 2.23E-153 | 4.72E-228 | 7.42E-92 | C5 |
| *S27698_1257* | CHH | 1.000 | 0.735 | 0.000 | 1.000 | 1.31E-41 | 5.33E-64 | 1.57E-220 | C6 |

| | | | | | | | | | |
|---|---|---|---|---|---|---|---|---|---|
| *S27698_1258* | CHH | 0.438 | 0.735 | 0.000 | 1.000 | 6.04E-11 | 5.33E-64 | 1.11E-159 | C6 |
| *S27698_1259* | CHH | 0.000 | 0.737 | 0.000 | 0.000 | 9.25E-36 | 2.29E-55 | 6.08E-19 | C1 |
| *S28865_10047* | CG | 0.000 | 0.011 | 1.000 | 1.000 | 1.00E+00 | 9.41E-147 | 1.00E+00 | WS |
| *S28865_10067* | CHG | 1.000 | 0.000 | 0.500 | 0.333 | 3.68E-06 | 1.00E+00 | 1.00E+00 | 5-Aza |
| *S28865_10080* | CHG | 0.498 | 0.000 | 1.000 | 1.000 | 1.00E+00 | 4.82E-07 | 1.00E+00 | WS |
| *S28865_10084* | CHH | 0.000 | 0.000 | 0.500 | 0.667 | 1.00E+00 | 3.89E-02 | 1.00E+00 | WS |
| *S28865_10091* | CHH | 0.000 | 0.000 | 0.500 | 0.667 | 1.00E+00 | 7.31E-04 | 1.00E+00 | WS |
| *S28865_10106* | CHH | 0.002 | 0.000 | 0.500 | 0.667 | 1.00E+00 | 6.54E-04 | 1.00E+00 | WS |
| *S28865_10108* | CG | 0.999 | 0.000 | 1.000 | 1.000 | 1.58E-251 | 8.83E-167 | 4.98E-43 | C5 |
| *S28865_10116* | CHH | 0.212 | 0.000 | 0.500 | 0.667 | 1.00E+00 | 3.42E-02 | 1.00E+00 | WS |
| *S28865_10186* | CHG | 0.214 | 0.996 | 1.000 | 1.000 | 4.85E-73 | 1.00E+00 | 1.13E-09 | C8 |
| *S28865_10190* | CHH | 0.214 | 1.000 | 0.500 | 0.333 | 2.80E-01 | 3.56E-02 | 2.91E-05 | C1 |
| *S29620_1172* | CG | 0.500 | 0.000 | 1.000 | 0.500 | 1.00E+00 | 1.07E-06 | 1.00E+00 | WS |
| *S29620_1182* | CHG | 0.000 | 0.000 | 0.000 | 0.500 | 1.00E+00 | 1.85E-07 | 1.00E+00 | WS |
| *S29620_1183* | CHH | 0.500 | 0.500 | 0.000 | 0.000 | 1.00E+00 | 5.04E-15 | 1.00E+00 | WS |
| *S29620_1206* | CHH | 1.000 | 0.500 | 1.000 | 0.000 | 1.00E+00 | 7.47E-18 | 1.00E+00 | WS |
| *S29620_1207* | CHH | 0.500 | 0.500 | 0.000 | 0.003 | 1.00E+00 | 8.03E-14 | 1.00E+00 | WS |
| *S29620_1213* | CHH | 0.500 | 0.500 | 0.000 | 0.000 | 1.00E+00 | 5.04E-15 | 1.00E+00 | WS |
| *S29620_1223* | CHG | 0.000 | 0.000 | 0.000 | 0.500 | 1.00E+00 | 2.11E-11 | 1.00E+00 | WS |
| *S29620_1225* | CG | 0.500 | 0.996 | 0.000 | 0.000 | 1.00E+00 | 6.29E-146 | 1.00E+00 | WS |
| *S29620_1281* | CHH | 0.500 | 0.500 | 1.000 | 1.000 | 1.00E+00 | 5.04E-15 | 1.00E+00 | WS |
| *S29620_1284* | CHH | 0.500 | 0.000 | 1.000 | 0.500 | 1.00E+00 | 5.04E-06 | 1.00E+00 | WS |
| *S41184_3999* | CG | 1.000 | 0.990 | 0.500 | 0.001 | 1.00E+00 | 4.94E-108 | 1.00E+00 | WS |
| *S41184_6323* | CHG | 0.000 | 1.000 | 0.000 | 0.000 | 2.91E-04 | 5.44E-04 | 1.00E+00 | WS |
| *S41184_6324* | CHH | 0.000 | 1.000 | 0.000 | 0.000 | 9.44E-04 | 1.77E-03 | 1.00E+00 | WS |
| *S41184_6329* | CHH | 0.000 | 1.000 | 0.000 | 0.000 | 9.44E-04 | 1.77E-03 | 1.00E+00 | WS |
| *S41184_6334* | CHH | 0.000 | 1.000 | 0.000 | 0.000 | 9.44E-04 | 1.77E-03 | 1.00E+00 | WS |
| *S41184_6337* | CHH | 0.000 | 1.000 | 0.000 | 0.000 | 9.44E-04 | 1.77E-03 | 1.00E+00 | WS |
| *S41184_6339* | CHH | 0.000 | 1.000 | 0.000 | 0.000 | 9.44E-04 | 1.77E-03 | 1.00E+00 | WS |
| *S41184_6347* | CHH | 0.000 | 1.000 | 0.000 | 0.000 | 9.44E-04 | 1.77E-03 | 1.00E+00 | WS |
| *S41184_6351* | CHH | 1.000 | 0.000 | 1.000 | 1.000 | 9.44E-04 | 1.77E-03 | 1.00E+00 | WS |

| | | | | | | | | | |
|---|---|---|---|---|---|---|---|---|---|
| *S41184_6364* | CHH | 0.000 | 1.000 | 0.000 | 0.000 | 9.44E-04 | 1.77E-03 | 1.00E+00 | WS |
| *S41184_6365* | CHH | 0.000 | 1.000 | 0.000 | 0.000 | 9.44E-04 | 1.77E-03 | 1.00E+00 | WS |
| *S44030_4857* | CHH | 0.012 | 0.617 | 0.000 | 0.000 | 2.87E-71 | 2.12E-74 | 1.32E-04 | C1 |
| *S44030_4865* | CG | 0.006 | 0.000 | 0.500 | 1.000 | 1.00E+00 | 2.55E-232 | 3.98E-20 | C3 |
| *S44030_4868* | CHG | 0.000 | 0.621 | 0.000 | 0.000 | 1.04E-91 | 9.99E-76 | 3.88E-07 | C1 |
| *S44030_4873* | CG | 0.000 | 0.000 | 0.000 | 1.000 | 1.00E+00 | 1.85E-232 | 3.03E-20 | C3 |
| *S44030_4876* | CHG | 0.000 | 0.000 | 0.000 | 1.000 | 1.00E+00 | 3.79E-232 | 4.57E-20 | C3 |
| *S44030_4879* | CHH | 0.000 | 0.621 | 0.000 | 0.004 | 3.38E-91 | 9.22E-67 | 1.33E-05 | C1 |
| *S44030_4888* | CHH | 0.003 | 0.621 | 0.000 | 1.000 | 8.85E-83 | 7.41E-38 | 1.00E+00 | 5-Aza |
| *S44030_4898* | CHH | 0.000 | 0.623 | 0.000 | 0.005 | 1.09E-91 | 5.84E-67 | 1.21E-05 | C1 |
| *S44030_4900* | CHH | 0.000 | 0.621 | 0.000 | 0.000 | 3.38E-91 | 3.24E-75 | 1.26E-06 | C1 |
| *S44030_4907* | CG | 1.000 | 0.000 | 0.500 | 1.000 | 2.69E-279 | 6.65E-232 | 4.06E-90 | C5 |
| *S44030_4910* | CHH | 0.000 | 0.621 | 0.000 | 0.000 | 3.38E-91 | 3.24E-75 | 1.26E-06 | C1 |
| *S44030_4915* | CG | 1.000 | 0.000 | 0.500 | 1.000 | 2.69E-279 | 6.65E-232 | 4.06E-90 | C5 |
| *S44030_4918* | CHG | 0.000 | 0.000 | 0.500 | 1.000 | 1.00E+00 | 2.74E-232 | 5.28E-18 | C3 |
| *S48520_2021* | CHG | 0.500 | 1.000 | 0.000 | 0.000 | 1.39E-03 | 1.18E-11 | 1.00E+00 | WS |
| *S48520_2022* | CHH | 0.000 | 1.000 | 0.000 | 0.001 | 7.66E-40 | 6.96E-45 | 1.59E-32 | C1 |
| *S48520_2032* | CG | 1.000 | 0.000 | 1.000 | 0.000 | 1.57E-40 | 1.00E+00 | 1.00E+00 | 5-Aza |
| *S48520_2042* | CG | 1.000 | 0.000 | 1.000 | 0.497 | 4.40E-03 | 1.00E+00 | 1.00E+00 | 5-Aza |
| *S48520_2055* | CG | 1.000 | 0.963 | 0.996 | 0.002 | 1.00E+00 | 7.11E-34 | 1.10E-23 | C7 |
| *S48520_2088* | CG | 1.000 | 0.000 | 1.000 | 1.000 | 1.57E-40 | 2.38E-47 | 1.51E-34 | C5 |
| *S48520_2122* | CG | 1.000 | 1.000 | 1.000 | 0.242 | 1.00E+00 | 2.58E-07 | 1.33E-01 | WS |
| *S48520_2122* | CG | 1.000 | 1.000 | 1.000 | 0.242 | 1.00E+00 | 2.58E-07 | 1.33E-01 | WS |
| *S48520_2125* | CHH | 0.000 | 0.964 | 0.000 | 0.002 | 5.83E-08 | 1.26E-34 | 1.81E-06 | C1 |
| *S48520_2125* | CHH | 0.000 | 0.964 | 0.000 | 0.002 | 5.83E-08 | 1.26E-34 | 1.81E-06 | C1 |
| *S48520_2137* | CHH | 0.000 | 1.000 | 0.004 | 0.759 | 3.88E-07 | 1.00E+00 | 1.00E+00 | 5-Aza |
| *S48520_2137* | CHH | 0.000 | 1.000 | 0.004 | 0.759 | 3.88E-07 | 1.00E+00 | 1.00E+00 | 5-Aza |
| *S53645_5199* | CG | 1.000 | 0.667 | 0.989 | 0.000 | 1.00E+00 | 5.38E-47 | 1.00E+00 | WS |
| *S53645_5218* | CHG | 1.000 | 0.333 | 0.000 | 0.000 | 1.00E+00 | 3.39E-46 | 1.00E+00 | WS |
| *S53645_5226* | CHH | 1.000 | 0.667 | 1.000 | 0.000 | 1.00E+00 | 2.62E-46 | 1.00E+00 | WS |
| *S53645_5228* | CHH | 1.000 | 0.333 | 0.000 | 0.000 | 1.00E+00 | 1.10E-45 | 1.00E+00 | WS |

| | | | | | | | | | |
|---|---|---|---|---|---|---|---|---|---|
| *S53645_5241* | CHH | 1.000 | 0.667 | 1.000 | 0.000 | 1.00E+00 | 2.62E-46 | 1.00E+00 | WS |
| *S53645_5252* | CHG | 1.000 | 0.667 | 1.000 | 0.000 | 1.00E+00 | 9.26E-46 | 1.00E+00 | WS |
| *S53645_5261* | CHH | 1.000 | 0.667 | 1.000 | 0.000 | 1.00E+00 | 2.62E-46 | 1.00E+00 | WS |
| *S53645_5267* | CHG | 1.000 | 0.667 | 0.996 | 0.000 | 1.00E+00 | 8.07E-47 | 1.00E+00 | WS |
| *S53645_5281* | CHG | 1.000 | 0.667 | 1.000 | 0.000 | 1.00E+00 | 9.26E-46 | 1.00E+00 | WS |
| *S53645_5285* | CHG | 1.000 | 0.333 | 0.000 | 0.000 | 1.00E+00 | 4.40E-46 | 1.00E+00 | WS |
| *S61275_25038* | CHH | 1.000 | 1.000 | 1.000 | 0.000 | 1.00E+00 | 3.36E-10 | 1.00E+00 | WS |
| *S61275_25075* | CHH | 0.000 | 0.000 | 0.000 | 0.500 | 1.00E+00 | 1.38E-10 | 1.00E+00 | WS |
| *S61275_25123* | CG | 0.750 | 1.000 | 1.000 | 0.000 | 1.00E+00 | 2.53E-11 | 1.68E-01 | WS |
| *S61275_25131* | CHH | 0.750 | 0.000 | 1.000 | 0.000 | 3.42E-23 | 1.00E+00 | 1.00E+00 | 5-Aza |
| *S61275_25135* | CHH | 0.500 | 0.250 | 1.000 | 0.000 | 1.00E+00 | 1.00E+00 | 2.21E-03 | C2 |
| *S61275_25149* | CHH | 0.000 | 0.250 | 1.000 | 0.500 | 1.00E+00 | 1.00E+00 | 3.42E-03 | C2 |
| *S61275_25170* | CHH | 0.000 | 0.500 | 1.000 | 0.500 | 2.19E-12 | 1.48E-04 | 1.78E-12 | C8 |
| *S61275_25171* | CHH | 0.000 | 0.500 | 1.000 | 0.000 | 2.19E-12 | 1.06E-05 | 2.17E-13 | C2 |
| *S61275_25176* | CHH | 0.000 | 0.000 | 1.000 | 0.500 | 1.00E+00 | 3.68E-09 | 1.00E+00 | WS |
| *S61275_25183* | CHG | 0.000 | 0.500 | 1.000 | 0.500 | 6.75E-13 | 1.00E+00 | 1.00E+00 | 5-Aza |
| *S61275_25194* | CHH | 0.000 | 0.500 | 1.000 | 0.500 | 1.00E+00 | 7.64E-05 | 1.00E+00 | WS |
| *S61275_25199* | CHH | 0.500 | 1.000 | 1.000 | 0.000 | 1.00E+00 | 8.03E-10 | 1.00E+00 | WS |
| *S61275_25218* | CHH | 0.000 | 0.000 | 0.000 | 0.500 | 1.00E+00 | 3.68E-09 | 1.00E+00 | WS |
| *S61275_25225* | CHH | 0.000 | 0.500 | 1.000 | 0.500 | 1.00E+00 | 7.64E-05 | 1.00E+00 | WS |
| *S62651_1962* | CHH | 0.000 | 1.000 | 1.000 | 0.000 | 2.11E-17 | 2.31E-15 | 3.87E-02 | C8 |
| *S62651_2001* | CHH | 0.999 | 0.000 | 0.000 | 0.500 | 1.98E-16 | 1.00E+00 | 1.00E+00 | 5-Aza |
| *S62651_2057* | CHH | 1.000 | 0.000 | 0.000 | 0.578 | 4.70E-14 | 1.00E+00 | 1.68E-20 | C4 |
| *S62651_2070* | CHH | 1.000 | 0.000 | 0.000 | 0.578 | 4.70E-14 | 1.00E+00 | 1.68E-20 | C4 |
| *S62651_2072* | CHH | 1.000 | 0.000 | 1.000 | 0.578 | 4.70E-14 | 1.00E+00 | 1.00E+00 | 5-Aza |
| *S62651_2073* | CHH | 0.993 | 0.000 | 1.000 | 0.583 | 1.26E-13 | 1.00E+00 | 1.00E+00 | 5-Aza |
| *S62651_2097* | CG | 1.000 | 1.000 | 1.000 | 0.500 | 1.00E+00 | 8.68E-06 | 1.93E-04 | C7 |
| *S62651_2106* | CHH | 1.000 | 0.000 | 0.000 | 0.500 | 6.54E-11 | 1.00E+00 | 1.73E-08 | C4 |
| *S62651_2107* | CHH | 1.000 | 0.000 | 0.000 | 0.500 | 6.54E-11 | 1.00E+00 | 1.70E-08 | C4 |
| *S62651_2128* | CG | 1.000 | 1.000 | 0.500 | 0.500 | 1.00E+00 | 8.49E-28 | 1.00E+00 | WS |
| *S63591_13851* | CHH | 1.000 | 0.500 | 0.000 | 0.000 | 3.61E-51 | 1.00E+00 | 2.60E-17 | C4 |

| | | | | | | | | | |
|---|---|---|---|---|---|---|---|---|---|
| *S63591_13855* | CHH | 1.000 | 0.500 | 0.000 | 0.000 | 3.61E-51 | 1.00E+00 | 2.60E-17 | C4 |
| *S63591_13858* | CHH | 1.000 | 0.500 | 0.000 | 0.000 | 3.61E-51 | 1.00E+00 | 2.60E-17 | C4 |
| *S63591_13860* | CHH | 1.000 | 0.500 | 0.000 | 0.000 | 3.61E-51 | 1.00E+00 | 2.60E-17 | C4 |
| *S63591_13869* | CHH | 1.000 | 0.500 | 0.000 | 0.000 | 3.82E-51 | 1.00E+00 | 2.63E-17 | C4 |
| *S63591_13932* | CHH | 1.000 | 0.569 | 0.000 | 0.500 | 7.77E-01 | 1.00E+00 | 1.89E-02 | C6 |
| *S63591_13934* | CHH | 1.000 | 0.567 | 0.000 | 0.502 | 2.71E-01 | 1.00E+00 | 4.18E-03 | C6 |
| *S63591_13935* | CHH | 1.000 | 0.569 | 0.000 | 0.500 | 7.77E-01 | 1.00E+00 | 1.89E-02 | C6 |
| *S63591_13955* | CHG | 0.003 | 0.430 | 1.000 | 0.957 | 7.49E-113 | 3.89E-03 | 9.15E-30 | C8 |
| *S63591_14014* | CG | 0.997 | 0.569 | 0.000 | 0.500 | 3.41E-01 | 1.00E+00 | 7.67E-03 | C6 |
| *S63591_15187* | CHH | 0.000 | 0.552 | 0.000 | 2.000 | 1.51E-53 | 1.33E-65 | 1.14E-24 | C1 |
| *S63591_15228* | CG | 0.000 | 0.448 | 0.000 | 0.000 | 1.20E-40 | 1.26E-52 | 8.96E-20 | C1 |
| *S63962_3472* | CG | 1.000 | 0.000 | 0.000 | 1.000 | 6.19E-50 | 1.25E-48 | 2.14E-03 | C6 |
| *S63962_3481* | CG | 1.000 | 0.000 | 1.000 | 1.000 | 6.43E-50 | 1.25E-48 | 1.00E+00 | 5-Aza |
| *S63962_3484* | CG | 1.000 | 0.000 | 1.000 | 1.000 | 6.19E-50 | 1.25E-48 | 1.00E+00 | 5-Aza |
| *S63962_3495* | CG | 1.000 | 0.000 | 1.000 | 1.000 | 6.19E-50 | 1.25E-48 | 1.00E+00 | 5-Aza |
| *S63962_3498* | CG | 1.000 | 0.000 | 1.000 | 1.000 | 6.19E-50 | 1.25E-48 | 1.00E+00 | 5-Aza |
| *S63962_3516* | CG | 0.997 | 0.000 | 1.000 | 1.000 | 2.90E-48 | 1.31E-48 | 1.00E+00 | 5-Aza |
| *S63962_3527* | CG | 0.999 | 0.000 | 1.000 | 1.000 | 1.04E-47 | 1.25E-48 | 1.00E+00 | 5-Aza |
| *S63962_3536* | CG | 0.994 | 0.032 | 1.000 | 0.000 | 2.24E-36 | 1.00E+00 | 1.00E+00 | 5-Aza |
| *S63962_3549* | CG | 0.997 | 0.000 | 1.000 | 1.000 | 2.90E-48 | 1.25E-48 | 1.00E+00 | 5-Aza |
| *S63962_3556* | CHG | 1.000 | 0.000 | 1.000 | 1.000 | 9.28E-50 | 1.87E-48 | 1.00E+00 | 5-Aza |
| *S64405_1933* | CHH | 0.000 | 0.000 | 0.000 | 0.987 | 1.00E+00 | 1.04E-151 | 2.03E-77 | C3 |
| *S64405_1936* | CG | 0.929 | 1.000 | 0.000 | 0.921 | 1.00E+00 | 3.35E-02 | 5.91E-38 | C6 |
| *S64405_1942* | CHG | 0.013 | 1.000 | 0.000 | 0.000 | 2.10E-119 | 7.87E-175 | 8.92E-78 | C1 |
| *S64405_1945* | CG | 0.987 | 1.000 | 0.000 | 0.993 | 1.00E+00 | 1.00E+00 | 1.72E-71 | C6 |
| *S64405_1952* | CG | 0.987 | 1.000 | 0.000 | 0.997 | 1.00E+00 | 1.00E+00 | 2.49E-74 | C6 |
| *S64405_1954* | CHG | 0.013 | 1.000 | 0.000 | 0.003 | 2.10E-119 | 4.89E-165 | 4.47E-73 | C1 |
| *S64405_1959* | CHG | 0.013 | 0.994 | 0.000 | 0.003 | 9.16E-110 | 1.27E-152 | 5.26E-67 | C1 |
| *S64405_1970* | CG | 0.994 | 0.000 | 0.000 | 1.000 | 3.88E-125 | 5.29E-175 | 0.00E+00 | C6 |
| *S64405_1976* | CG | 1.000 | 0.994 | 0.001 | 1.000 | 1.00E+00 | 1.00E+00 | 1.81E-92 | C6 |
| *S64405_2002* | CHH | 0.000 | 1.000 | 0.000 | 0.000 | 6.57E-135 | 1.71E-174 | 7.32E-88 | C1 |

| | | | | | | | | | |
|---|---|---|---|---|---|---|---|---|---|
| *S64428_4014* | CG | 0.995 | 0.000 | 0.992 | 0.000 | 1.60E-22 | 1.00E+00 | 1.00E+00 | 5-Aza |
| *S64428_4015* | CHG | 1.000 | 0.000 | 0.992 | 0.000 | 3.09E-24 | 1.00E+00 | 1.00E+00 | 5-Aza |
| *S64428_4020* | CG | 1.000 | 0.000 | 1.000 | 0.995 | 2.05E-24 | 2.43E-23 | 4.94E-20 | C5 |
| *S64428_4025* | CHH | 0.566 | 1.000 | 0.991 | 0.000 | 1.67E-01 | 7.04E-25 | 1.25E-43 | C7 |
| *S64428_4031* | CG | 0.437 | 1.000 | 1.000 | 0.997 | 1.53E-04 | 1.00E+00 | 2.46E-04 | C8 |
| *S64428_4058* | CHH | 0.000 | 1.000 | 0.000 | 0.000 | 1.00E+00 | 3.91E-23 | 1.00E+00 | WS |
| *S64428_4073* | CHG | 0.000 | 1.000 | 1.000 | 0.000 | 1.57E-21 | 2.17E-25 | 2.47E-90 | C8 |
| *S64428_4077* | CHG | 1.000 | 0.000 | 1.000 | 0.000 | 1.57E-21 | 1.00E+00 | 1.00E+00 | 5-Aza |
| *S64428_4081* | CHH | 0.000 | 1.000 | 0.333 | 0.000 | 5.10E-21 | 6.98E-25 | 1.84E-19 | C1 |
| *S64428_4086* | CG | 0.012 | 1.000 | 1.000 | 1.000 | 4.52E-19 | 1.00E+00 | 7.44E-17 | C8 |
| *S64428_4087* | CHG | 1.000 | 0.000 | 0.488 | 0.000 | 1.57E-21 | 1.00E+00 | 1.00E+00 | 5-Aza |
| *S64428_4090* | CHG | 0.000 | 0.000 | 0.488 | 0.000 | 1.00E+00 | 1.00E+00 | 3.34E-18 | C2 |
| *S64428_4094* | CHH | 0.000 | 1.000 | 0.000 | 0.000 | 5.10E-21 | 6.98E-25 | 8.74E-19 | C1 |
| *S64428_4105* | CHG | 1.000 | 0.000 | 0.012 | 0.997 | 1.57E-21 | 6.09E-24 | 1.28E-79 | C6 |
| *S64428_4108* | CHG | 0.000 | 1.000 | 0.988 | 1.000 | 1.57E-21 | 1.00E+00 | 3.99E-15 | C8 |
| *S64428_4120* | CHH | 0.000 | 1.000 | 0.000 | 1.000 | 5.10E-21 | 1.00E+00 | 1.00E+00 | 5-Aza |
| *S6490_14847* | CG | 0.001 | 0.000 | 0.243 | 0.000 | 1.00E+00 | 1.00E+00 | 2.36E-23 | C2 |
| *S6490_14896* | CHH | 0.000 | 0.000 | 0.501 | 0.500 | 1.00E+00 | 1.36E-07 | 1.00E+00 | NA |
| *S6490_14904* | CHH | 0.001 | 0.003 | 0.742 | 0.500 | 1.00E+00 | 2.51E-84 | 3.63E-09 | C2 |
| *S6490_14905* | CHH | 0.001 | 0.000 | 0.742 | 0.500 | 1.00E+00 | 2.01E-89 | 9.25E-11 | C2 |
| *S6490_14910* | CHH | 0.001 | 0.000 | 0.485 | 0.500 | 1.00E+00 | 5.10E-122 | 1.37E-20 | C2 |
| *S6490_14914* | CHG | 0.001 | 0.000 | 0.482 | 0.000 | 1.00E+00 | 1.00E+00 | 8.86E-16 | C2 |
| *S6490_14923* | CHH | 0.001 | 0.000 | 0.482 | 0.500 | 1.00E+00 | 4.15E-122 | 9.31E-21 | C2 |
| *S6490_14925* | CHH | 0.001 | 0.000 | 0.490 | 0.000 | 1.00E+00 | 1.00E+00 | 1.04E-15 | C2 |
| *S6490_14929* | CG | 0.001 | 0.000 | 0.485 | 0.000 | 1.00E+00 | 1.00E+00 | 4.29E-16 | C2 |
| *S6490_14931* | CG | 0.004 | 0.000 | 0.485 | 0.000 | 1.00E+00 | 1.00E+00 | 3.45E-14 | C2 |
| *S6490_14948* | CHH | 0.001 | 0.003 | 1.000 | 1.000 | 1.00E+00 | 5.50E-117 | 1.00E+00 | WS |
| *S6490_14950* | CHH | 0.001 | 0.000 | 0.485 | 0.500 | 1.00E+00 | 4.15E-122 | 1.30E-20 | C2 |
| *S6490_14951* | CHH | 0.001 | 0.000 | 0.485 | 0.500 | 1.00E+00 | 4.15E-122 | 1.30E-20 | C2 |
| *S66538_3446* | CG | 1.000 | 0.214 | 1.000 | 1.000 | 2.85E-66 | 5.02E-129 | 2.82E-03 | C5 |
| *S66538_3450* | CHH | 1.000 | 0.214 | 1.000 | 1.000 | 1.38E-65 | 2.42E-128 | 1.40E-02 | C5 |

| | | | | | | | | | |
|---|---|---|---|---|---|---|---|---|---|
| *S66538_3452* | CHH | 1.000 | 0.214 | 1.000 | 1.000 | 1.38E-65 | 2.42E-128 | 1.40E-02 | C5 |
| *S66538_3454* | CHH | 0.362 | 0.000 | 1.000 | 1.000 | 1.72E-08 | 7.66E-135 | 1.00E+00 | WS |
| *S66538_3455* | CHH | 1.000 | 0.214 | 1.000 | 0.000 | 8.21E-65 | 1.00E+00 | 1.00E+00 | 5-Aza |
| *S66538_3463* | CHH | 0.627 | 0.214 | 1.000 | 1.000 | 1.46E-19 | 2.42E-128 | 1.00E+00 | WS |
| *S66538_3466* | CHH | 1.000 | 0.214 | 0.000 | 1.000 | 1.38E-65 | 2.42E-128 | 1.28E-15 | C6 |
| *S66538_3472* | CG | 0.627 | 0.500 | 1.000 | 0.995 | 3.08E-20 | 1.70E-118 | 2.33E-12 | C5 |
| *S66538_3474* | CHH | 0.627 | 0.000 | 1.000 | 1.000 | 1.78E-21 | 6.48E-133 | 4.74E-15 | C5 |
| *S66538_3483* | CHH | 0.000 | 0.000 | 0.500 | 0.000 | 1.00E+00 | 1.00E+00 | 3.63E-51 | C2 |
| *S66538_3492* | CHH | 0.627 | 0.000 | 0.000 | 1.000 | 1.78E-21 | 1.15E-110 | 4.55E-126 | C3 |
| *S66538_3494* | CHH | 0.627 | 0.000 | 0.000 | 1.000 | 1.78E-21 | 1.15E-110 | 5.42E-126 | C3 |
| *S66538_3503* | CHH | 0.627 | 0.000 | 0.000 | 0.000 | 1.78E-21 | 1.00E+00 | 1.44E-14 | C4 |
| *S66538_3648* | CHH | 0.000 | 0.500 | 0.000 | 0.000 | 6.49E-09 | 1.34E-12 | 6.09E-04 | C1 |
| *S6731_11973* | CHH | 0.333 | 0.024 | 0.999 | 0.333 | 1.00E+00 | 1.00E+00 | 1.43E-140 | C2 |
| *S6731_11985* | CHH | 0.333 | 0.024 | 1.000 | 0.667 | 1.00E+00 | 6.67E-192 | 1.00E+00 | WS |
| *S6731_12004* | CHH | 0.333 | 0.524 | 0.997 | 0.333 | 1.00E+00 | 1.00E+00 | 4.73E-09 | C8 |
| *S6731_12012* | CHH | 0.333 | 0.025 | 1.000 | 0.333 | 1.00E+00 | 1.00E+00 | 5.55E-144 | C2 |
| *S6731_12019* | CHG | 0.333 | 0.264 | 1.000 | 0.667 | 1.00E+00 | 1.21E-09 | 1.00E+00 | WS |
| *S6731_12036* | CHG | 0.500 | 0.000 | 0.995 | 0.667 | 1.00E+00 | 3.61E-200 | 1.00E+00 | WS |
| *S6731_12077* | CHH | 0.000 | 0.000 | 1.000 | 0.500 | 1.00E+00 | 2.66E-188 | 1.00E+00 | WS |
| *S6731_12080* | CHH | 0.000 | 0.000 | 1.000 | 0.500 | 1.00E+00 | 2.66E-188 | 1.00E+00 | WS |
| *S6731_12106* | CHH | 0.000 | 0.000 | 1.000 | 0.002 | 1.00E+00 | 1.00E+00 | 5.56E-84 | C2 |
| *S6731_12107* | CHH | 0.000 | 4.000 | 0.997 | 0.000 | 1.00E+00 | 1.00E+00 | 1.12E-90 | C2 |
| *S6731_12108* | CHH | 0.000 | 0.000 | 1.000 | 0.000 | 1.00E+00 | 1.00E+00 | 5.51E-89 | C2 |
| *S6731_12123* | CG | 0.500 | 0.957 | 0.997 | 0.500 | 8.58E-139 | 1.92E-185 | 0.00E+00 | C8 |
| *S6731_12129* | CHH | 0.003 | 0.534 | 1.000 | 0.000 | 5.09E-40 | 8.46E-65 | 5.83E-222 | C2 |
| *S6731_12132* | CHH | 0.000 | 0.488 | 0.000 | 0.000 | 1.47E-41 | 3.91E-57 | 9.99E-21 | C1 |
| *S6731_12133* | CHH | 0.000 | 0.483 | 0.000 | 0.000 | 3.81E-41 | 1.29E-56 | 1.75E-20 | C1 |
| *S6731_12134* | CHH | 0.000 | 0.483 | 0.000 | 0.000 | 6.02E-41 | 1.74E-56 | 2.17E-20 | C1 |
| *S6731_12146* | CHH | 0.000 | 0.486 | 0.000 | 0.000 | 2.15E-41 | 8.13E-57 | 1.35E-20 | C1 |
| *S6731_12148* | CHH | 0.000 | 0.474 | 0.000 | 0.000 | 5.21E-40 | 1.00E+00 | 1.00E+00 | 5-Aza |
| *S71193_2245* | CHH | 0.000 | 1.000 | 0.333 | 0.667 | 2.11E-13 | 3.68E-35 | 1.32E-23 | C8 |

| | | | | | | | | | |
|---|---|---|---|---|---|---|---|---|---|
| *S71193_2248* | CG | 0.267 | 1.000 | 1.000 | 1.000 | 6.92E-08 | 1.00E+00 | 1.03E-01 | 5-Aza |
| *S71193_2257* | CHH | 0.000 | 0.500 | 0.000 | 0.333 | 6.25E-19 | 7.40E-99 | 7.40E-10 | C1 |
| *S71193_2263* | CHH | 0.000 | 0.500 | 0.000 | 0.333 | 6.25E-19 | 7.40E-99 | 7.40E-10 | C1 |
| *S71193_2268* | CHG | 0.000 | 1.000 | 0.500 | 0.667 | 1.20E-19 | 1.00E+00 | 1.00E+00 | 5-Aza |
| *S71193_2271* | CHH | 0.733 | 1.000 | 0.038 | 0.333 | 1.00E+00 | 1.37E-102 | 1.27E-01 | WS |
| *S71193_2276* | CHH | 0.267 | 0.500 | 25.000 | 0.333 | 1.30E-06 | 7.40E-99 | 4.73E-10 | C1 |
| *S71193_2286* | CHH | 0.733 | 1.000 | 25.000 | 0.333 | 1.00E+00 | 6.77E-102 | 1.29E-01 | WS |
| *S71193_2296* | CHH | 0.267 | 0.500 | 0.358 | 0.333 | 1.00E+00 | 3.11E-16 | 1.54E-05 | C1 |
| *S71193_2299* | CHH | 0.733 | 0.500 | 1.000 | 0.667 | 1.30E-06 | 7.40E-99 | 5.36E-05 | C5 |
| *S71193_2306* | CG | 1.000 | 0.000 | 1.000 | 0.665 | 1.60E-20 | 1.08E-94 | 1.88E-15 | C5 |
| *S71193_2308* | CG | 1.000 | 1.000 | 0.667 | 1.000 | 1.00E+00 | 1.00E+00 | 7.61E-16 | C6 |
| *S71193_2311* | CHG | 0.000 | 0.000 | 0.667 | 0.667 | 1.00E+00 | 4.11E-102 | 1.00E+00 | WS |
| *S71193_2330* | CHH | 1.000 | 1.000 | 0.642 | 0.667 | 1.00E+00 | 2.32E-101 | 9.59E-12 | C7 |
| *S71193_2342* | CHH | 0.267 | 1.000 | 0.463 | 0.500 | 2.86E-07 | 1.00E+00 | 1.00E+00 | 5-Aza |
| *S71193_2357* | CHH | 0.000 | 0.500 | 0.465 | 0.500 | 1.87E-02 | 1.00E+00 | 1.00E+00 | 5-Aza |
| *S71479_2341* | CG | 0.667 | 0.000 | 0.000 | 0.000 | 2.26E-20 | 1.00E+00 | 8.75E-15 | C4 |
| *S71479_2411* | CHH | 0.000 | 1.000 | 0.000 | 0.000 | 1.83E-35 | 2.38E-55 | 3.69E-31 | C1 |
| *S71479_2455* | CHH | 0.000 | 0.000 | 0.000 | 0.500 | 1.00E+00 | 5.59E-53 | 6.91E-33 | C3 |
| *S71479_2470* | CHH | 0.000 | 0.000 | 0.000 | 0.500 | 1.00E+00 | 5.59E-53 | 7.00E-33 | C3 |
| *S71479_2474* | CHH | 0.000 | 0.000 | 0.000 | 0.500 | 1.00E+00 | 5.59E-53 | 7.00E-33 | C3 |
| *S71479_2504* | CHH | 0.000 | 0.000 | 0.500 | 1.000 | 1.00E+00 | 2.09E-53 | 1.55E-27 | C3 |
| *S71479_2509* | CHH | 0.000 | 0.000 | 0.500 | 1.000 | 1.00E+00 | 2.39E-53 | 2.50E-26 | C3 |
| *S71479_2511* | CHH | 0.000 | 0.000 | 0.500 | 1.000 | 1.00E+00 | 2.24E-53 | 2.46E-26 | C3 |
| *S71479_2515* | CHG | 0.000 | 0.000 | 0.500 | 0.992 | 1.00E+00 | 2.61E-47 | 1.00E+00 | WS |
| *S71479_2522* | CHH | 0.000 | 0.000 | 0.500 | 1.000 | 1.00E+00 | 2.09E-53 | 2.41E-26 | C3 |
| *S72001_4230* | CHG | 0.499 | 0.000 | 0.000 | 0.000 | 2.69E-12 | 1.00E+00 | 1.00E+00 | 5-Aza |
| *S72001_4231* | CHH | 0.498 | 0.000 | 0.000 | 0.000 | 2.09E-11 | 1.00E+00 | 1.00E+00 | 5-Aza |
| *S72001_4233* | CHH | 0.499 | 0.000 | 0.000 | 0.000 | 8.72E-12 | 1.00E+00 | 1.00E+00 | 5-Aza |
| *S72001_4235* | CG | 0.500 | 0.000 | 0.000 | 0.000 | 3.59E-13 | 1.00E+00 | 1.00E+00 | 5-Aza |
| *S72001_4236* | CHG | 0.500 | 0.000 | 0.000 | 0.000 | 5.43E-13 | 1.00E+00 | 1.00E+00 | 5-Aza |
| *S72001_4244* | CHH | 0.500 | 0.000 | 0.000 | 0.000 | 1.76E-12 | 1.00E+00 | 1.00E+00 | 5-Aza |

| | | | | | | | | | |
|---|---|---|---|---|---|---|---|---|---|
| *S72001_4245* | CHH | 0.500 | 0.111 | 0.002 | 0.000 | 2.03E-07 | 1.00E+00 | 1.00E+00 | 5-Aza |
| *S72001_4253* | CHH | 0.499 | 0.000 | 0.002 | 0.000 | 8.81E-12 | 1.00E+00 | 1.00E+00 | 5-Aza |
| *S72001_4254* | CHH | 0.498 | 0.000 | 0.002 | 0.000 | 2.08E-11 | 1.00E+00 | 1.00E+00 | 5-Aza |
| *S72001_4367* | CHH | 0.999 | 0.000 | 1.000 | 1.000 | 7.55E-12 | 1.00E+00 | 1.00E+00 | 5-Aza |
| *S76299_8902* | CHH | 1.000 | 0.500 | 0.000 | 0.000 | 3.05E-118 | 1.00E+00 | 1.00E+00 | 5-Aza |
| *S76299_8946* | CG | 1.000 | 0.824 | 1.000 | 1.000 | 1.76E-16 | 1.00E+00 | 1.00E+00 | 5-Aza |
| *S76299_8963* | CG | 1.000 | 0.676 | 0.500 | 0.000 | 3.18E-40 | 1.00E+00 | 1.00E+00 | 5-Aza |
| *S76299_8973* | CHH | 0.000 | 0.322 | 0.500 | 1.000 | 3.42E-39 | 1.00E+00 | 1.00E+00 | 5-Aza |
| *S76299_8995* | CHH | 1.000 | 1.000 | 0.000 | 1.000 | 1.00E+00 | 1.00E+00 | 4.58E-03 | C6 |
| *S76299_8996* | CHH | 1.000 | 0.333 | 0.000 | 0.000 | 2.87E-79 | 1.00E+00 | 4.40E-29 | C4 |
| *S76299_9003* | CHH | 0.000 | 0.333 | 0.000 | 1.000 | 1.00E+00 | 3.62E-103 | 2.97E-29 | C3 |
| *S76299_9004* | CHH | 1.000 | 0.667 | 0.000 | 1.000 | 5.49E-76 | 3.12E-101 | 1.33E-129 | C6 |
| *S76299_9005* | CHH | 1.000 | 0.333 | 0.000 | 0.000 | 2.87E-79 | 1.00E+00 | 4.40E-29 | C4 |
| *S76299_9018* | CHH | 1.000 | 0.500 | 0.000 | 0.500 | 1.77E-33 | 1.94E-34 | 2.47E-52 | C6 |
| *S76299_9020* | CHH | 0.989 | 0.500 | 0.000 | 0.500 | 1.20E-29 | 2.29E-34 | 2.74E-49 | C6 |
| *S76299_9022* | CHH | 1.000 | 0.500 | 0.000 | 0.500 | 2.62E-33 | 2.58E-34 | 3.74E-52 | C6 |
| *S76299_9029* | CHH | 1.000 | 0.250 | 0.000 | 0.000 | 1.83E-127 | 1.00E+00 | 3.19E-36 | C4 |
| *S76299_9031* | CHH | 1.000 | 0.500 | 0.000 | 0.500 | 1.95E-33 | 2.18E-34 | 2.80E-52 | C6 |
| *S76299_9032* | CHH | 1.000 | 0.250 | 0.000 | 0.000 | 1.59E-127 | 1.00E+00 | 3.14E-36 | C4 |
| *S76299_9095* | CHH | 1.000 | 0.250 | 0.000 | 0.000 | 1.59E-127 | 1.00E+00 | 3.14E-36 | C4 |
| *S76299_9105* | CHG | 1.000 | 0.338 | 0.000 | 0.000 | 8.10E-03 | 1.00E+00 | 1.00E+00 | 5-Aza |
| *S76299_9110* | CHG | 1.000 | 0.500 | 0.000 | 0.497 | 1.28E-37 | 1.93E-36 | 1.03E-55 | C6 |
| *S76981_6017* | CHH | 0.000 | 0.000 | 0.333 | 0.000 | 1.00E+00 | 1.00E+00 | 7.13E-10 | C2 |
| *S76981_6018* | CHH | 0.000 | 0.000 | 0.333 | 0.000 | 1.00E+00 | 1.00E+00 | 7.11E-10 | C2 |
| *S76981_6034* | CHH | 0.000 | 0.000 | 0.333 | 0.000 | 1.00E+00 | 1.00E+00 | 7.11E-10 | C2 |
| *S76981_6046* | CHH | 0.000 | 0.486 | 0.000 | 0.000 | 2.24E-60 | 2.75E-89 | 7.25E-14 | C1 |
| *S76981_6047* | CHH | 0.000 | 0.493 | 0.333 | 0.000 | 7.76E-65 | 9.94E-96 | 3.59E-55 | C1 |
| *S76981_6050* | CHH | 0.000 | 0.487 | 0.333 | 0.000 | 8.03E-62 | 9.41E-93 | 1.51E-53 | C1 |
| *S76981_6053* | CHH | 0.000 | 0.494 | 0.000 | 0.000 | 1.42E-14 | 4.26E-85 | 1.00E+00 | WS |
| *S76981_6054* | CHH | 0.000 | 0.500 | 0.000 | 0.000 | 4.85E-16 | 1.03E-95 | 1.00E+00 | WS |
| *S76981_6057* | CHG | 0.000 | 0.500 | 0.000 | 0.000 | 1.49E-16 | 3.18E-96 | 1.00E+00 | WS |

| | | | | | | | | | |
|---|---|---|---|---|---|---|---|---|---|
| *S76981_6063* | CHH | 0.000 | 0.987 | 0.000 | 0.000 | 3.53E-68 | 7.50E-83 | 1.00E+00 | WS |
| *S76981_6095* | CHG | 1.000 | 0.013 | 1.000 | 1.000 | 1.09E-68 | 2.31E-83 | 1.00E+00 | WS |
| *S77906_11496* | CHG | 1.000 | 0.829 | 1.000 | 1.000 | 2.43E-02 | 1.01E-08 | 1.00E+00 | WS |
| *S77906_11502* | CHG | 0.000 | 0.700 | 0.298 | 0.662 | 1.78E-35 | 6.03E-25 | 1.06E-19 | C1 |
| *S77906_11522* | CHH | 0.000 | 0.842 | 0.000 | 0.592 | 1.11E-13 | 6.60E-08 | 1.00E+00 | 5-Aza |
| *S77906_11533* | CHG | 1.000 | 1.000 | 1.000 | 0.746 | 1.00E+00 | 1.15E-17 | 1.14E-03 | C7 |
| *S77906_11557* | CHH | 1.000 | 0.829 | 0.018 | 0.929 | 7.90E-02 | 1.00E+00 | 5.85E-26 | C6 |
| *S77906_14506* | CG | 0.000 | 1.000 | 0.000 | 0.000 | 8.12E-50 | 1.22E-264 | 5.83E-42 | C1 |
| *S77906_14526* | CG | 0.000 | 1.000 | 0.000 | 0.000 | 8.12E-50 | 1.22E-264 | 5.83E-42 | C1 |
| *S77906_14551* | CG | 0.000 | 1.000 | 0.591 | 0.005 | 6.74E-06 | 7.35E-07 | 8.87E-15 | C1 |
| *S77906_14562* | CG | 0.000 | 0.997 | 0.167 | 1.000 | 4.88E-47 | 1.00E+00 | 1.00E+00 | 5-Aza |
| *S77906_14593* | CG | 0.000 | 1.000 | 0.002 | 0.000 | 8.12E-50 | 1.22E-264 | 2.55E-43 | C1 |
| *S77915_4573* | CHG | 1.000 | 0.500 | 1.000 | 1.000 | 1.00E+00 | 2.60E-46 | 1.00E+00 | WS |
| *S77915_4595* | CHH | 1.000 | 0.500 | 0.000 | 0.000 | 1.00E+00 | 8.42E-46 | 1.00E+00 | WS |
| *S77915_4604* | CHH | 1.000 | 0.500 | 0.000 | 0.000 | 1.00E+00 | 8.42E-46 | 1.00E+00 | WS |
| *S77915_4612* | CHH | 1.000 | 0.500 | 0.000 | 0.000 | 1.00E+00 | 8.42E-46 | 1.00E+00 | WS |
| *S77915_4629* | CHG | 1.000 | 0.500 | 1.000 | 1.000 | 1.00E+00 | 2.60E-46 | 1.00E+00 | WS |
| *S77915_4639* | CHH | 1.000 | 0.500 | 0.000 | 0.000 | 1.00E+00 | 8.42E-46 | 1.00E+00 | WS |
| *S77915_4657* | CHH | 1.000 | 0.500 | 0.000 | 0.000 | 1.00E+00 | 8.42E-46 | 1.00E+00 | WS |
| *S77915_4674* | CHH | 0.036 | 1.000 | 1.000 | 1.000 | 2.22E-26 | 1.00E+00 | 1.00E+00 | 5-Aza |
| *S77915_4780* | CG | 1.000 | 0.500 | 0.000 | 0.000 | 1.00E+00 | 1.73E-46 | 1.00E+00 | WS |
| *S77915_4782* | CHH | 0.012 | 0.500 | 1.000 | 1.000 | 1.00E+00 | 8.42E-46 | 1.00E+00 | WS |
| *S77915_4787* | CHG | 1.000 | 0.000 | 0.000 | 0.000 | 5.60E-53 | 1.00E+00 | 6.14E-01 | 5-Aza |
| *S77915_4802* | CHH | 0.012 | 0.500 | 1.000 | 1.000 | 1.00E+00 | 8.42E-46 | 1.00E+00 | WS |
| *S77915_4803* | CHH | 0.012 | 0.500 | 1.000 | 1.000 | 1.00E+00 | 8.42E-46 | 1.00E+00 | WS |
| *S77915_4806* | CHH | 0.012 | 0.500 | 1.000 | 1.000 | 1.00E+00 | 1.02E-45 | 1.00E+00 | WS |
| *S77915_4807* | CHH | 0.015 | 0.500 | 1.000 | 1.000 | 1.00E+00 | 1.12E-45 | 1.00E+00 | WS |
| *S77915_4811* | CHH | 0.012 | 0.500 | 1.000 | 1.000 | 1.00E+00 | 8.42E-46 | 1.00E+00 | WS |
| *S77915_4817* | CG | 1.000 | 0.500 | 1.000 | 1.000 | 7.23E-51 | 1.73E-46 | 4.28E-03 | C5 |
| *S77915_4840* | CG | 1.000 | 0.500 | 1.000 | 1.000 | 7.23E-51 | 1.73E-46 | 4.28E-03 | C5 |
| *S77915_4857* | CG | 1.000 | 0.000 | 0.000 | 0.000 | 3.73E-53 | 1.00E+00 | 4.01E-01 | 5-Aza |

| | | | | | | | | | |
|---|---|---|---|---|---|---|---|---|---|
| *S8686_2487* | CG | 1.000 | 1.000 | 1.000 | 0.011 | 1.00E+00 | 4.55E-82 | 1.00E+00 | WS |
| *S8686_2496* | CHH | 0.000 | 0.000 | 0.000 | 0.989 | 1.00E+00 | 2.76E-80 | 1.00E+00 | WS |
| *S8686_2506* | CHH | 0.000 | 0.000 | 0.000 | 0.989 | 1.00E+00 | 7.75E-81 | 1.00E+00 | WS |
| *S8686_2514* | CHH | 0.000 | 0.000 | 0.000 | 0.989 | 1.00E+00 | 2.20E-81 | 1.00E+00 | WS |
| *S8686_2515* | CHH | 0.000 | 0.000 | 0.000 | 0.989 | 1.00E+00 | 3.64E-79 | 1.00E+00 | WS |
| *S8686_2521* | CHH | 0.000 | 0.000 | 0.000 | 0.978 | 1.00E+00 | 6.07E-76 | 1.00E+00 | WS |
| *S8686_2522* | CHH | 0.000 | 0.000 | 0.000 | 0.989 | 1.00E+00 | 2.20E-81 | 1.00E+00 | WS |
| *S8686_2523* | CHH | 0.000 | 0.000 | 0.000 | 0.989 | 1.00E+00 | 1.50E-80 | 1.00E+00 | WS |
| *S8686_2536* | CHG | 1.000 | 0.000 | 0.500 | 0.989 | 4.13E-05 | 6.79E-82 | 6.45E-05 | C5 |
| *S8686_2561* | CHH | 1.000 | 0.000 | 0.000 | 0.989 | 1.34E-04 | 2.20E-81 | 1.81E-09 | C6 |
| *S8686_2575* | CHH | 0.000 | 0.000 | 0.000 | 0.989 | 1.00E+00 | 2.20E-81 | 1.00E+00 | WS |
| *S8686_2576* | CHG | 1.000 | 0.000 | 1.000 | 0.000 | 4.13E-05 | 1.00E+00 | 1.00E+00 | 5-Aza |
| *S8686_2587* | CG | 0.000 | 1.000 | 0.500 | 1.000 | 2.74E-05 | 1.00E+00 | 1.00E+00 | 5-Aza |
| *S9768_1644* | CHG | 1.000 | 1.000 | 1.000 | 0.222 | 1.00E+00 | 1.05E-72 | 1.00E+00 | WS |
| *S9768_1655* | CHG | 1.000 | 0.000 | 0.000 | 1.000 | 4.28E-40 | 2.23E-83 | 6.28E-03 | C6 |
| *S9768_1671* | CHH | 1.000 | 0.000 | 0.000 | 0.000 | 1.39E-39 | 1.00E+00 | 6.72E-24 | C4 |
| *S9768_1672* | CHH | 0.000 | 0.000 | 0.000 | 0.778 | 1.00E+00 | 3.40E-72 | 3.98E-21 | C3 |
| *S9768_1682* | CHG | 0.000 | 1.000 | 0.990 | 1.000 | 4.28E-40 | 1.00E+00 | 2.02E-20 | C8 |
| *S9768_1710* | CHH | 0.000 | 1.000 | 1.000 | 0.278 | 1.39E-39 | 2.07E-70 | 8.14E-95 | C8 |
| *S9768_1728* | CHH | 0.000 | 0.000 | 0.667 | 0.000 | 1.00E+00 | 1.00E+00 | 3.49E-21 | C2 |
| *S9768_1731* | CG | 1.000 | 1.000 | 0.333 | 0.000 | 1.00E+00 | 1.49E-83 | 1.00E+00 | WS |
| *S9768_1735* | CHH | 0.000 | 1.000 | 1.000 | 0.278 | 1.39E-39 | 2.07E-70 | 8.14E-95 | C8 |
| *S9768_1747* | CHH | 0.000 | 0.000 | 0.667 | 0.278 | 1.00E+00 | 1.00E+00 | 1.74E-17 | C2 |
| *S9768_1767* | CHG | 0.000 | 1.000 | 1.000 | 0.500 | 3.26E-05 | 1.07E-01 | 5.95E-10 | C8 |

8

**Fig. S1.** Effects of maternal genotype ($F_1$-mother identity) on global methylation and cytosines contexts. Note that each family consist in four plants, each one assigned to one of the four levels of the 2x2 treatment combination design.

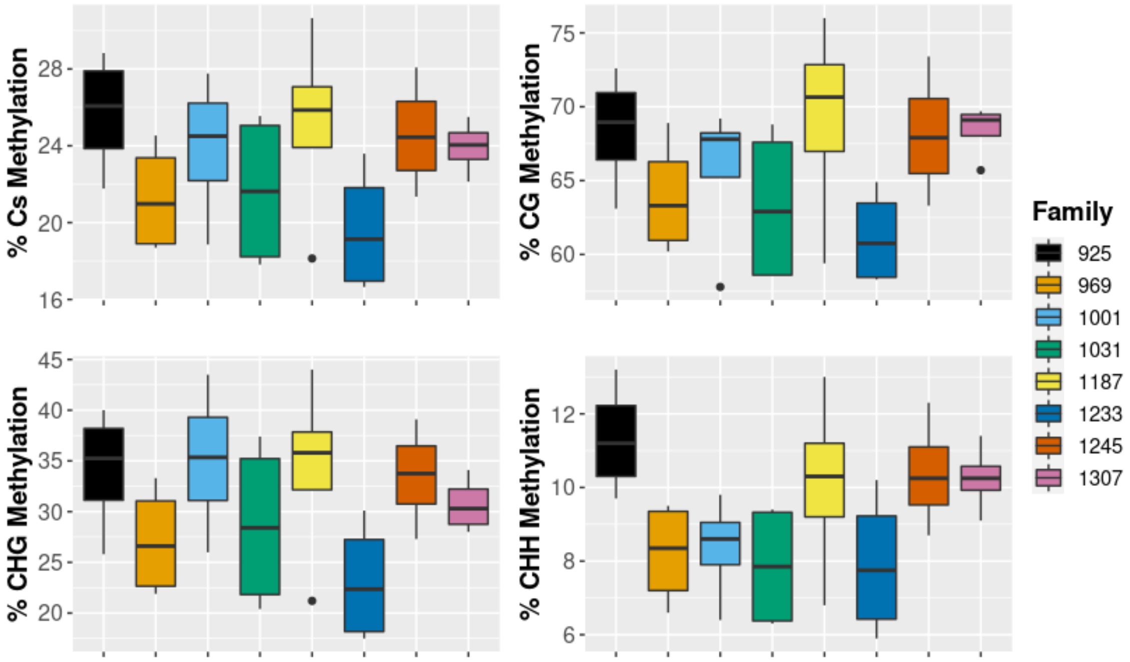

**Fig. S2.** Distribution of DMCs on scaffolds of the *E. cicutarium* genome. Most DMCs appeared on different scaffolds (mean DMCs per scaffold 1.65 ± 0.03).

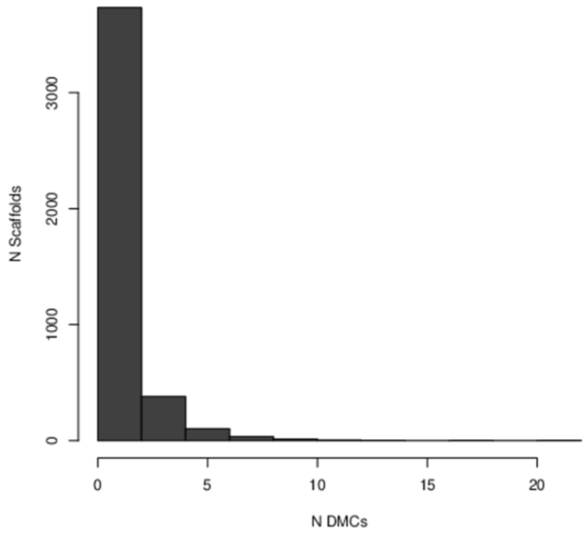

**Fig. S3.** Plot of within-groups sum of squares against number of clusters using the K-means algorithm using the average methylation level of the DMCs for each experimental group. After 8 clusters (in red), the observed difference in the within-cluster dissimilarity is despicable.

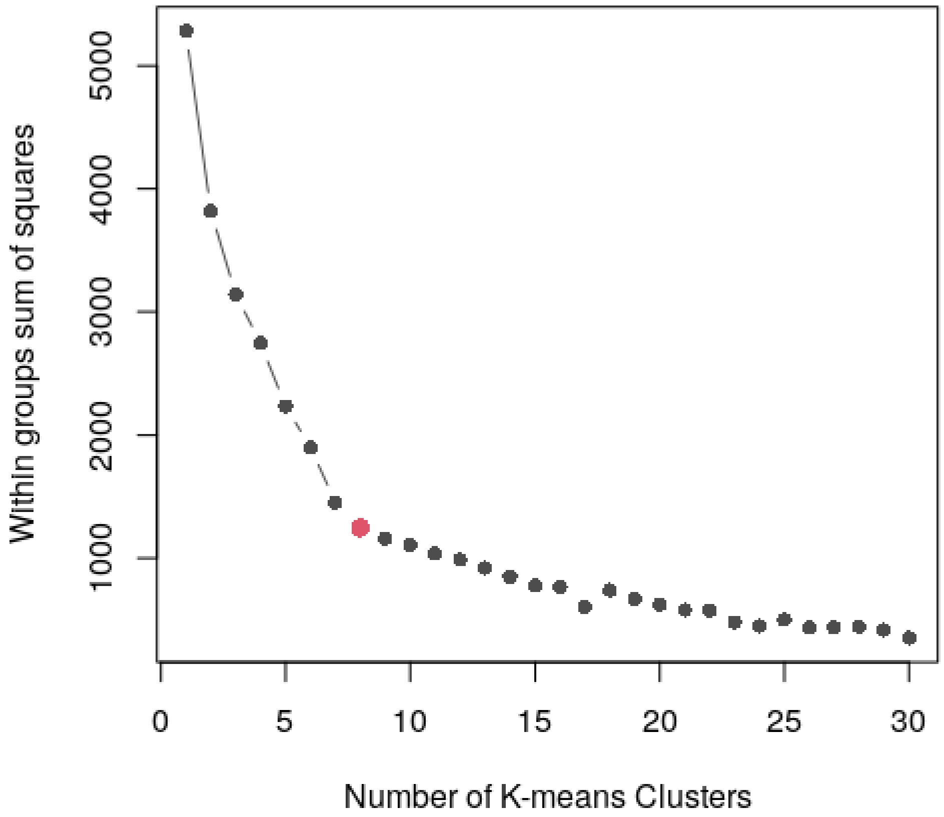

**Fig. S4.** Distribution of DMCs overlapping transposable elements at different regions.

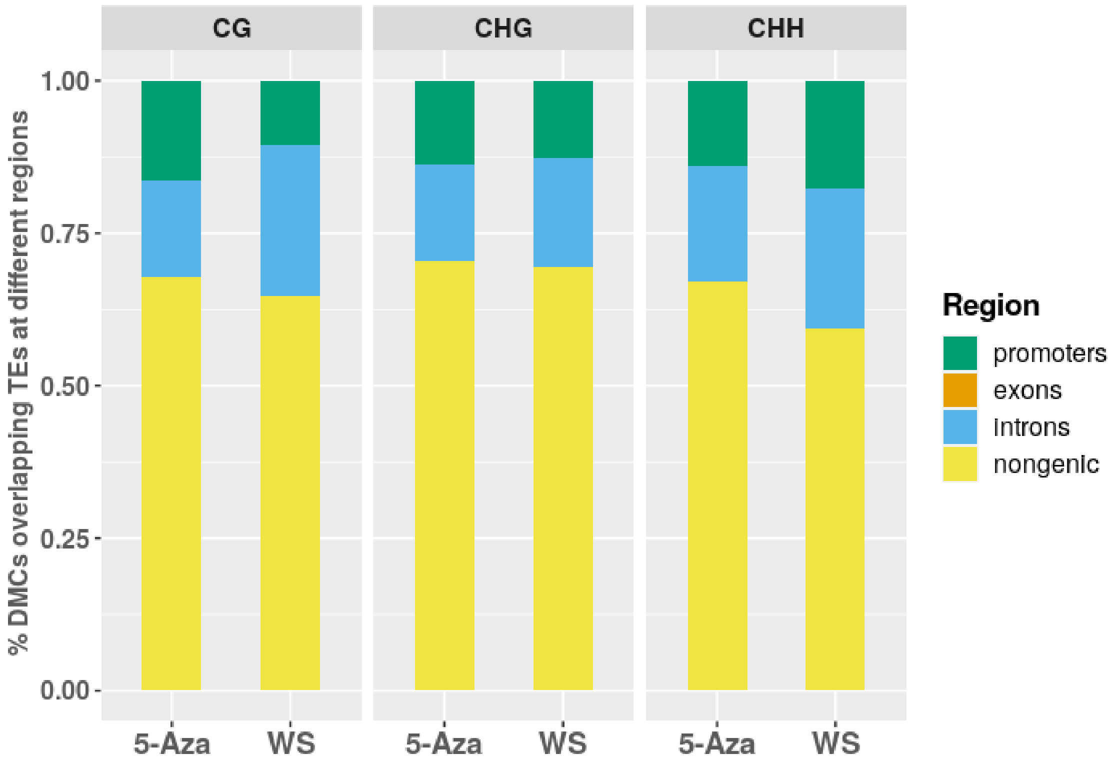
